# Linking Kinetochore Attachment to Checkpoint Control: The Role of Aurora B in BubR1 Acetylation

**DOI:** 10.1101/2025.08.22.671725

**Authors:** Si-Young Choi, Hyungmin Kim, Sung-Soo Kim, Sanghyo Park, Hyunsook Lee

## Abstract

We report here that Aurora B kinase is critical in BubR1 acetylation at K250, transducing kinetochore attachment status to spindle assembly checkpoint (SAC) activity. Aurora B phosphorylates BubR1 at serines 16 and 39 in an unattachment-dependent manner, which is a prerequisite for BubR1 acetylation at lysine 250 (K250). Utilizing a novel anti-AcK250 monoclonal antibody in structured illumination microscopy (SIM), we reveal that BubR1 acetylation is associated with kinetochore expansion, evidenced by crescent-shaped volume increase of ZW10 and Mad2 surrounding the kinetochore. Furthermore, K250-BubR1 acetylation enhances its interaction with CENP-E, a kinesin crucial for chromosome alignment through facilitating end-on attachment. Disrupting Aurora B-mediated phosphorylation of Ser16/39 impaired K250 acetylation, compromising both fibrous corona polymerization and mitotic checkpoint complex (MCC) maintenance. Conversely, introducing K250 acetylation mimetic form rescued MCC and fibrous corona in cells expressing phosphorylation-deficient S16A/S39A mutants. Our findings reveal Aurora B-BubR1 acetylation axis as a critical SAC signaling pathway, interlinking unattachment sensing with kinetochore expansion and SAC activity. We propose that this stepwise phosphorylation-acetylation on BubR1 constitutes a crucial part of SAC signaling code, transducing unattachment signal to fibrous corona polymerization and SAC activity. This phosphorylation-acetylation cascade in BubR1 offers potential therapeutic targets in treating cancers of chromosome instability.

## INTRODUCTION

The integrity of chromosomes is ensured by the precise segregation of the replicated genome into two daughter cells, a process that critically relies on elegant coordination between chromosome-spindle attachment and the spindle assembly checkpoint (SAC). The SAC is activated at kinetochores lacking end-on microtubule attachments, arresting mitosis until all kinetochores achieve amphitelic attachment (1–4). However, how mechanical status of kinetochore-microtubule (KT-MT) attachments relays to the SAC is not fully elucidated.

The formation and maintenance of the mitotic checkpoint complex (MCC), composed of BubR1, Mad2, Bub3, and CDC20, are central in SAC activity. Simultaneous with MCC binding to APC/C, fibrous corona polymerizes resulting in kinetochore expansion, increasing the chance of end-on KT-MT attachment. When all kinetochores are end-on attached, anaphase begins with the disassembly of MCC, which leads to the activation of APC/C-Cdc20 (5–10). A central question is how mechanical signal of kinetochore attachment status is synchronized with SAC activity.

Our previous research demonstrated that acetylation of BubR1 at lysine 250 (lysine 243 in mice) modulates APC/C activity and is pivotal for spindle assembly checkpoint signaling (11). This acetylation, which occurs exclusively during prometaphase, requires the tumor suppressor BRCA2, which acts as a scaffold facilitating PCAF acetyltransferase to associate with BubR1. Disruption of the BRCA2-BubR1 interaction in mice leads to tumorigenesis characterized by chromosome instability (12), mirroring the phenotype observed in BRCA2-deficient mice (13,14). Consistently, a heterozygous acetylation-deficient K243R mutation (equivalent to K250 in human) in *BubR1* (*K243R/+*) induces tumorigenesis, manifested by defects in error correction, weakened SAC, and ultimate chromosome mis-segregation (15). These studies indicated that BubR1 acetylation at K250 has dual roles: controlling KT-MT attachment and maintaining SAC integrity.

During mitosis, kinetochores initially attach to the lateral sides of microtubules. This lateral attachment is subsequently converted to end-on attachments, where microtubules directly connect to the outer kinetochore (16). The fibrous corona, a transient structure at unattached kinetochores, expands to enhance the probability of kinetochore-microtubule (KT-MT) interactions (17,18) and to facilitate the recruitment of the Mad1-Mad2 complex (3,19,20). At this stage, kinetochores are laterally attached, and SAC activation is linked to the dynamic volume expansion of outer kinetochore components (21,22).

CENP-E (Centromere Protein E), a plus-ended kinesin-like motor protein, is a component of the outer kinetochore that transports chromosomes along existing microtubules toward the spindle equator (23), supporting lateral attachment and facilitating the conversion to stable endon microtubule attachments at the fibrous corona. Concurrently, the corona’s building block, the RZZ (ROD, ZW10, Zwilch) complex, associates with dynein, a minus-ended motor, via the adaptor Spindly (24,25). The microtubule motors with opposite polarities, CENP-E and dynein-dynactin, promote biorientation from the kinetochore. Once end-on KT-MT attachment is achieved, the SAC is satisfied (26), and the fibrous corona disentangles, stabilizing the endon attachments (17,19,21,22,27). This volumetric regulation of fibrous corona is associated with SAC activity. However, the coordinated molecular mechanisms controlling unattachment sensing, kinetochore expansion, and SAC activation remain elusive.

Aurora B kinase (AurkB) is part of the chromosomal passenger complex (CPC) that localizes to the inner kinetochore. By phosphorylating the KMN network (Knl1, Mis12, Ndc80), the core microtubule-binding site at the outermost kinetochore (28) that recruits RZZ complex and Mad1-Mad2 (29) as well, AurkB destabilizes mis-attached kinetochores (30). Upon achieving correct bi-polar attachment, phosphatases neutralize AurkB’s activity, leading to the stabilization of KT-MT attachments and anaphase begins (31,32).

BubR1 associates with CENP-E (33). Notably, BubR1 acetylation at K250 affects its binding to CENP-E: the acetylation-deficient form lacks the binding, while the acetylation-mimetic form binding to CENP-E increases (15). Therefore, it is possible that AurkB activity is involved in BubR1 acetylation and CENP-E-assisted chromosome congression. However, how AurkB and BubR1 acetylation is connected has remained elusive.

Here, we demonstrate that AurkB phosphorylates BubR1 at serines 16 and 39, which is essential for BubR1 acetylation at K250 upon unattachment. BubR1 acetylation is associated with the polymerization of fibrous corona, simultaneously enhancing the binding to CENP-E and MCC maintenance. Collectively, AurkB-BubR1 acetylation axis constitutes an essential part of SAC signaling that interlinks unattachment signal to kinetochore expansion, control of end-on attachment, and MCC maintenance.

## MATERIAL AND METHODS

### Statistical Analysis

All Graphs and statistical analyses were generated using GraphPad Prism 5 software. Unless otherwise specified, *p-*values were calculated using a Student’s *t*-test, with results expressed as mean ± standard error of the mean (s.e.m.), where applicable.

### siRNAs and Plasmids

The following siRNAs were employed in this study: siLuciferase (Bioneer), siBRCA2 (5’- GAAGAACAAUAUCCUACUA-3’) (12), siBubR1-3’UTR (5’-GUCUCACAGAUUGCUGC CU-3’)(11), siAuroraB (5’-AAGAGCCUGUCACCCCAUCUG-3’), siAPC15 (5’-GUCUGGU CUAAGUUUCUUU-3’), and siCENP-E (5’-AAGGCUACAAUGGUACUAUAU).

mCherry-tagged BubR1 variants-including *S16A, S39A, S49A, S361A, T600A, S619A, S649A, T654A, T493A;S495A, T508A;S509A, S435A. S16A;K250R, S39A;K250R, S16A;S39A, S16D, S39D, S16D;K250R*, and *S39D;K250R*-were cloned into the pcDNA3.1-mCherry vector (34) for immunofluorescence, immunoprecipitation assay followed by Western blotting (IP-WB), and live-cell imaging experiments.

Full-length and truncated BubR1 constructs (amino acids 1-150, 1-322, 1-514, 322-1050, 514-760, and 760-1050) were cloned into the pcDNA3.1-myc vector (11) to identify the Aurora B-interacting region of BubR1 by IP-WB. Aurora B wild-type and its kinase-dead mutant (*K106R*) were cloned into the pcDNA3.1-myc vector for subsequent assays. Each construct was validated by sequencing, and expression was confirmed by Western blotting prior to experimental use.

### Cell culture and transfection

All cells used in this study were cultured in Dulbecco’s Modified Eagle’s Medium (DMEM; Serana) supplemented with 10% fetal bovine serum (FBS; Serana), 100U/mL penicillin, and 100μg/mL streptomycin (Penicillin/Streptomycin; Lonza). Cultures were maintained at 37°C in a humidified incubator with 5% CO_2_. Transfections of siRNAs and plasmids were carried out using Lipofectamine 2000 transfection reagent (Invitrogen) according to the manufacturer’s instructions. Unless otherwise noted, cells were incubated for 48 hours post-transfection before being used in subsequent experiments.

### Antibodies

The monoclonal antibody detecting acetylated BubR1 (anti-AcK250 mAb) was developed in in-house (34,35). Commercially sourced antibodies used in this study include: anti-BubR1 (612503; BD Biosciences), anti-α-tubulin-FITC (F2168; Upstate), anti-Centromere protein antibody (15-235; Antibodies inc.), anti-HDAC2 (ab12169; Abcam), anti-HDAC3 (ab7030; Abcam), anti-CDC20 (A15656; ABclonal), anti-Mad2 (A11469; ABclonal), anti-Cyclin B (sc-245; Santa Cruz Biotechnology), anti-β-actin (A700-057; Bethyl), anti-CDC27 (BD 610455; BD Biosciences), anti-FLAG (F1804; Sigma-Aldrich), anti-GAPDH (#2118; Cell Signaling Technology), anti-CENP-A (ab13939;Abcam), anti-ZW10 (sc-81430; Santa Cruz Biotechnology), anti-Myc (sc-40; Santa Cruz Biotechnology), anti-AIM-1 (BD611082; BD Biosciences), anti-Vinculin (sc-73614; Santa Cruz Biotechnology), anti-AurB-pT232 (636102; BioLegend), anti-Plk1 (ab17057; Abcam), anti-mCherry (ab125096; Abcam), anti-Bub3 (611730; BD Biosciences), ani-CENP-E (A15263; ABclonal), Cyclin B (sc-245; Santa Cruz Biotechnology), anti-pH3 (06-570; EMD Millipore), anti-pan-Phospho-serine (AP0932; ABclonal), and anti-Cdc25C (A12234, ABclonal).

### Cell cycle synchronization and drug treatments

To synchronize cells in mitosis, the following compounds were used for 20 hours: nocodazole (200ng/mL, M1404; Sigma), Paclitaxel/Taxol (2μM, T7402; Sigma), Monastrol (100μM, S8439; Selleckchem), and MG132 (10μM, 474790; Sigma). For mitotic chromosomes spread, colcemid (0.1mg/mL, 15212012; Gibco) was used.

For inducible expression of BubR1 in HeLa-FRT-TO cells, doxycycline (1μg/mL, 10592-13-9; Sigma) was administered for 48 hours. For kinase inhibition assays, the following inhibitors were added during the final 2 hours of a 20-hour nocodazole treatment: BI2536 (10nM, S1109; Selleckchem), Reversine (2μM, R3904; Sigma), Hesperadin (200nM, S1529; Selleckchem), and ZM447439 (2μM, S110301; Selleckchem). To inhibit HDAC activity, trichostatin A (10μM, T5882; Sigma) was used.

### Fluorescence microscopy and Structured Illumination Microscopy (SIM)

Conventional fluorescence images were acquired using a CoolSNAP HQ2 cooled CCD camera mounted on a DeltaVision Spectris Restoration microscope (GE Healthcare Life Sciences), equipped with a 60× oil-immersion objective lens (numerical aperture 1.42). Image deconvolution was performed using the SoftWorx software suite provided by GE Healthcare Life Sciences.

Structured illumination microscopy (SIM) was conducted using an Applied Precision DeltaVision OMX system (OMX Ultra High-Resolution Fluorescence Microscope, in SNU CMCI), equipped with a 60x oil-immersion objective (numerical aperture 1.42, refractive index 1.518, Olympus). SIM image reconstruction was performed using SoftWorx software with default settings (k_0_ angles: -0.810000, Wiener filter constant: 0.0030, bias offset: 35.00), along with channel-specific optical transfer functions (OTFs), k_0_ angles, and Wiener filters. Reconstructed images were aligned in the Z-axis using the BGR image source drawer.

### 3D reconstruction and volume measurement

To classify kinetochores based on microtubule attachment status-no attachment, single attachment, or bi-attachment-Z-stack images were acquired at 0.125μm intervals using structured illumination microscopy. Image stacks were processed using Image J Fiji. Preprocessing included application of a Gaussian blur filter (raidus: 1.00 pixel) to reduce high-frequency noise, followed by background subtraction using the Subtract Background function (rolling ball radius: 5 pixels) to enhance signal-to-noise ratio. For three-dimensional reconstruction, selected kinetochores were visualized using the 3D projection tool in imageJ Fiji. The brightest-point projection method was used, with X-axis rotation and interpolation enabled, to maintain structural integrity across Z-planes. The attachment status was inferred by analyzing the spatial patterns of CREST and microtubule signals. Fluorescent signal volumes corresponding to acetylated BubR1 (AcK250), total BubR1, Mad2, and ZW10 were quantified using the 3D Objects Counter plugin. Threshold values were determined empirically for each channel and condition. As an example, a voxel intensity threshold 20-300 (8-bit scale) was typically used to detect AcK250 signals, while minimizing non-specific background. Only objects falling within a defined volume range (0.01-3.0 μm^3^) and overlapping with CREST were included in the quantification. Volume data were collected from multiple cells across at least two independent experiments. Only objects within a defined volume range and signal intensity threshold were considered for quantitative analysis to ensure consistency and reduce variability. All volume measurements were exported and further analyzed using GraphPad Prism. Statistical comparisons between groups were conducted using non-parametric Mann-Whitney U test or Student’s *t*-test.

### Immunofluorescence assay

Fixation was done with 4% formaldehyde, prepared by diluting a 16% formaldehyde solution (methanol-free, Thermo Scientific), and then permeabilized with Tris-buffered saline (TBS) containing 0.5% Triton X-100 (TBS-0.5% TX). The cells were subsequently incubated with primary antibodies in an antibody dilution buffer (TBS-0.1% TX, 0.1% sodium azide, and 2% BSA) for overnight. After washing with TBS-0.1% TX, secondary antibodies were applied for 2 hours. Following washing with TBS-0.1% TX, mounting was done with Vectashield antifade mounting medium (Vector laboratories).

### Mitotic chromosomes spread

Cells were treated with 0.1 mg/ml of colcemid for 18 hours at 37 °C to arrest them in mitosis, followed by a brief washout for 5 minutes. The cells were then subjected to swelling in a hypotonic solution (0.075M of KCl) before being fixed with a methanol/acetic acid mixture (3:1). Finally, the cells were dropped onto slides for subsequent immunofluorescent staining.

### Mass spectrometry

HeLa-FRT-TO cells expressing *BubR1-WT*, *-K250R*, or *-K250Q* were cultured in 150-mm dishes. Ectopic expression of BubR1 was induced by the addition of doxycycline (1μg/mL) for 3 days. To arrest cells in mitosis, nocodazole (200ng/mL) was added for the last 16 hours. Cells were collected by mitotic shake-off, and protein lysates were prepared in lysis buffer containing 150mM NaCl, 0.5% NP-40, 1mM EDTA, 20mM Tris-Cl (pH 8.0), and protease inhibitor cocktail (Roche). The lysates were sonicated and incubated overnight with ANTI-FLAG M2 Affinity Gel (Sigma). After incubation, the beads were washed with lysis buffer, and the bound proteins were eluted. Protein concentrations were determined, and the eluates were separated by SDS-PAGE and stained using silver staining. Parallel samples were analyzed by Western blotting with anti-BubR1 and anti-FLAG antibodies. Gel sections containing the target bands were excised and sent for LC-MS analysis.

Sample preparation and proteomic analysis were performed by Bertis. The excised gels were destained with a solution of 50% acetonitrile (ACN, J.T. Baker) in 25 mM ammonium bicarbonate (ABC, sigma), then dehydrated with 100% ACN. Proteins were reduced with 10 mM dithiothreitol (DTT, Sigma) at 56°C for 30 minutes, then alkylated with 25 mM iodoacetamide (IAA, Sigma) at RT in the dark for 30 minutes. Proteolytic digestion was performed with trypsin at a 25:1 protein-to-trypsin ratio (w/w) at 37°C for 16 hours. Peptides were extracted from the gel in 20%, 50%, and 99% ACN solutions containing 0.1% trifluoroacetic acid (TFA, Thermo Fisher). Extracted peptides were dried, reconstituted in 0.1% TFA. For mass spectrometric analysis, tryptic peptides were reconstituted in 0.1% TFA and loaded onto a C18 analytical column (75 μm ID × 50 cm, PepMap RSLC C18, Thermo Fisher Scientific) for separation. Peptides were eluted with a linear gradient of 5-40% ACN in 0.1% TFA. The HPLC system was coupled to an Orbitrap Exploris 480 (Thermo Fisher Scientific) operating in Data-Dependent Acquisition (DDA) mode. The DDA analysis used the UniProt database (May 2022 release), with manually curated and *BubR1-K250R* and *BubR1-K250Q* variant sequences. Data analysis was performed with Proteome Discoverer software (version 2.4.1.15). Carbamidomethylation of cysteine (+ 57.021 Da) was set as a fixed modification, while oxidation of methionine (+ 15.995 Da) and phosphorylation of serine, threonine, and tyrosine (+ 79.9799 Da) were considered variable modifications.

### IP-WB analysis

Cells were harvested and resuspended in lysis buffer (150mM NaCl, 0.5% NP-40, 1mM EDTA, 20mM Tris-Cl pH 8.0) with protease inhibitor cocktail tablets (Complete Mini, EDTA-free, Roche). One milligram of protein was subjected to immunoprecipitation using a specific antibody, and protein G conjugated to Sepharose (Protein G Sepharose 4 Fast Flow, Merck). 50μg of protein was used to detect the expression level of proteins in total cell lysates (TCL).

### In vitro phosphorylation assay

MBP tagged-truncated BubR1 (amino acids 1-150) constructs-wild-type (WT), S39A, S16A;S39A, and S16A-were transformed into *E.coli* Rosetta strain. A single colony was inoculated into 3mL of LB medium and cultured overnight (∼18 hours), then transferred to 400mL of fresh medium and grown until the optical density at 600nm (OD_600_) reached 0.6∼0.7. Peptide expression was induced with 0.1mM isopropyl b-D-1-thiogalactopyranoside (IPTG, C-8001-1;Bioneer). Cells were harvested lysed in lysis buffer containing 50mM Tris-Cl pH 7.5, 2mM β-mercaptoethanol, 5mM imidazole, 5% glycerol, 0.5% NP-40, 0.1mM lysozyme, 1.5mg/mL DNase 1, 1mM PMSF, and EDTA-free protease inhibitor cocktail (Roche). Lysates were incubated with Ni-NTA agarose resin (1018244; QIAGEN) for 1 hour. Eluted proteins were concentrated using Amicon Ultra centrifugal filters (30kDa cutoff, UFC9030; Millipore) and further purified by size exclusion chromatography. For in vitro translation of Myc-tagged BubR1 constructs, TnT Quick coupled transcription/translation system (L1170; Promega) was used according to the manufacturer’s instructions.

For in vitro phosphorylation assay, either *E.coli*-purified or in vitro translated truncated BubR1 peptides were incubated with Aurora B kinase. Aurora B was either immunoprecipitated from mitotic or interphase cell lysates (obtained by mitotic shake-off), or obtained as a recombinant protein (14-835; Millipore). Reactions were carried out in phosphorylation buffer containing 10mM HEPES pH 7.5, 20mM KCl, 5mM MgCl_2_, 1mM DTT, and 100μM ATP. For detection of phosphorylation, either 5μCi of [γ-^32^P] ATP was added to the reaction mixture for autoradiography-based detection, or a phospho-specific antibody was used for immunoblotting.

### Live cell imaging

Time-lapse live cell imaging was performed using 60x/numerical aperture 1.42 lens (GE Healthcare Life Sciences) on a DeltaVision microscope (GE Healthcare Life Sciences) on CO_2_ chamber at 37°C as previously described(11,15). Cells were cultured in a glass-bottom dish (8-well chamber, Lab-Tek II Chambered Coverglass, Thermo Fisher Sci.) and analyzed 24 hours post-transfection. Time-lapse images were captured every 5 minutes, deconvolved and maximally projected using SoftWoRx software.

## RESULTS

### Lysine 250 acetylation of BubR1 identifies unattached kinetochore

BubR1 K250 is acetylated exclusively during prometaphase, and this specific modification dictates whether BubR1 acts as an inhibitor or a substrate of APC/C during metaphase-anaphase transition (11). Acetylated BubR1 (AcK250) is a component of MCC (mitotic checkpoint complex) (11), while deacetylation of AcK250 triggers MCC disassembly due to BubR1 degradation (15).

Notably, loss of BubR1 acetylation in mice (*K243R/+*) leads to spontaneous tumorigenesis due to chromosome mis-segregation, accompanied by aberrant KT-MT attachments and premature MCC disassembly (15). These observations indicate that K250-BubR1 acetylation is involved in both stable KT-MT interactions and MCC maintenance, extending the understanding of the established role of BubR1 in linking KT-MT attachment to SAC activity (36,37). However, the precise mechanism coordinating K250 acetylation/deacetylation with the mode of kinetochore attachment remained unclear.

To investigate how BubR1 acetylation responds to KT-MT attachment status, we developed a novel monoclonal antibody that specifically targets acetylated BubR1 (anti-AcK250 mAb, Supplementary Fig. S1A & B) (34), applied it in a high-resolution immunofluorescence (IFA) technique, then explored how BubR1 acetylation responds to the mode of KT-MT attachment. Analysis of metaphase chromosome spreads, prepared after colcemid treatment and a brief 5-minute release, coupled with IFA confirmed that acetylated BubR1 was present at unattached kinetochores (Fig. 1A). In detail, there were three distinct IFA patterns: chromosomes with AcK250-BubR1 at only one sister kinetochore (Fig. 1A and B, Single); kinetochores lacking AcK250-BubR1 (Fig. 1A and B, No); and paired sister kinetochores with AcK250-BubR1 (Fig. 1A and B, Paired). These patterns suggested a possibility that AcK250-BubR1 might discriminate between unattached and attached kinetochores.

**Figure 1.**
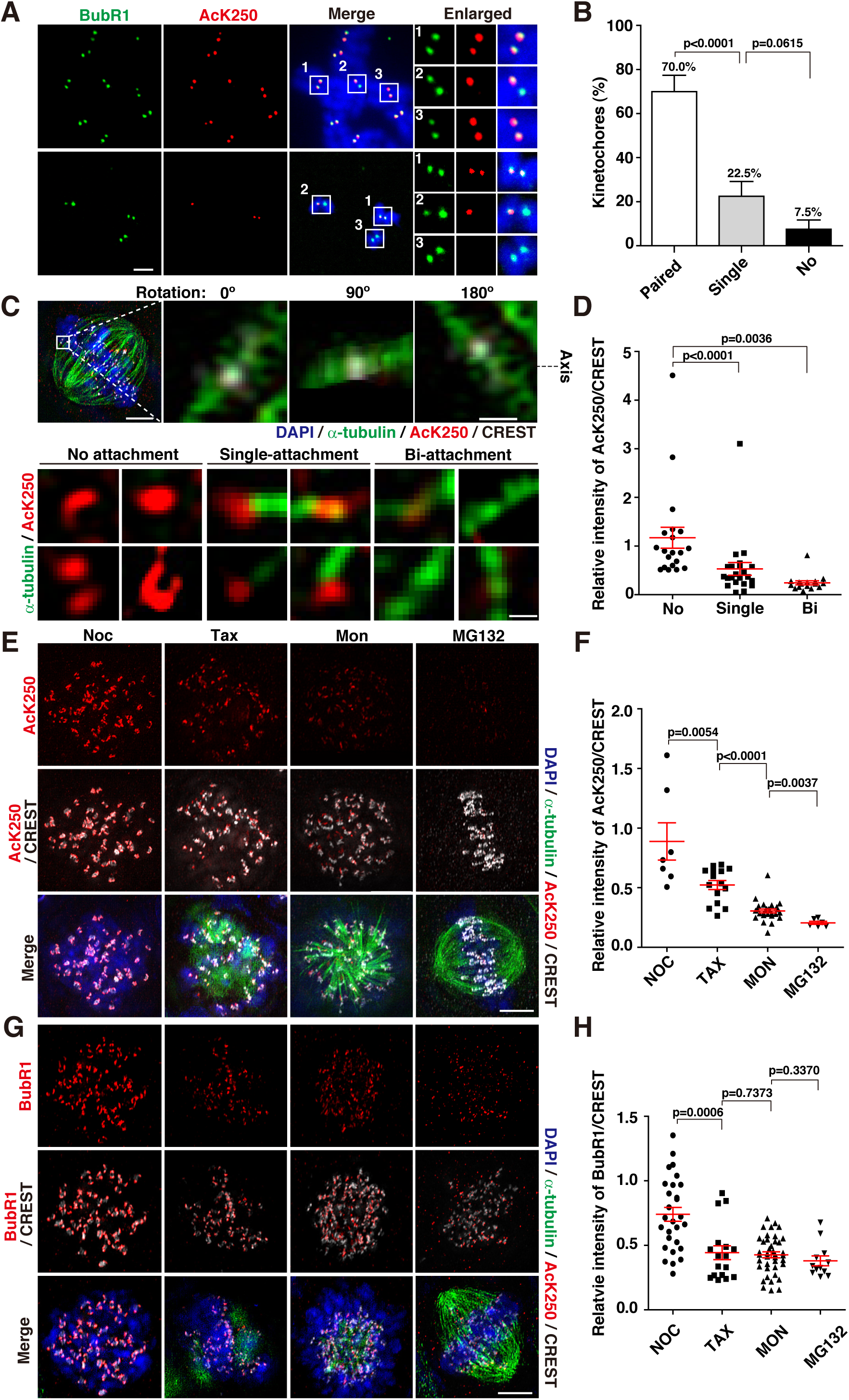
Acetylation of BubR1 at K250 identifies unattached kinetochore. (A) Immunofluorescent staining of metaphase chromosome spreads. Mitotic HeLa cells were enriched by treatment with colcemid (100 ng/mL) for 18 h, followed by a 5-minute release, and then subjected to co-immunostaining using anti-BubR1 and anti-acetylated BubR1 (AcK250) monoclonal antibodies (mAbs). Kinetochores were categorized into three groups based on anti-AcK250 mAb immunostaining: those immunostained at both sister kinetochores (Paired); one kinetochore (Single); or neither (No). Scale bar, 5 μm. (B) Proportion of anti-AcK250 mAb immunostaining patterns in (A). Results are from two independent experiments (mean ± s.e.m.; n = 40 chromosomes). P values were obtained using a t-test. (C) Representative images of microtubule attachment at kinetochores using structured illumination microscopy (SIM). Mitotic HeLa cells were treated with 10 μM MG132 to enrich the metaphase population, fixed, and then co-immunostained using anti-α-tubulin, anti-AcK250, and CREST antibodies. CREST-marked kinetochores were divided into three groups based on 3D-reconstructed images: unattached kinetochore (No attachment), monotelic kinetochore (Single-attachment), and amphitelic kinetochore (Bi-attachment). Scale bar, 5 μm (Top). Bottom, 3D-reconstructed images of anti-AcK250 mAb staining. Scale bar, 0.5 μm. The SIM images were acquired using an AP DeltaVision OMX Ultra High-Resolution Fluorescence Microscope (GE Healthcare). Image reconstruction and alignment were performed using SoftWoRx software. Three-dimensional reconstruction was performed using ImageJ software (3D projection). (D) Relative intensity of anti-AcK250 immunostaining normalized to CREST. Number of kinetochores scored: No attachment (No), n = 20; monotelic attachment (Single), n = 22; bipolar attachment (Bi), n = 16. The graphs represent two independent experiments (mean ± s.e.m.). Statistical differences between groups were assessed using the Mann-Whitney U test; p values are indicated. (E-H) HeLa cells treated with 200 ng/mL nocodazole (Noc), 2 μM paclitaxel/taxol (Tax), 100 μM Monastrol (Mon), or 10 μM MG132, respectively, were co-immunostained with CREST, anti-α-tubulin, and anti-AcK250 (E, F) or anti-BubR1 (G, H). (F, H) Graphs are from two independent experiments (mean±s.e.m.). Scale bar, 5 μm.

To assess whether AcK250-BubR1 specifically identifies unattached kinetochores, growing HeLa cells were treated with the proteasome inhibitor MG132 to enrich cells in metaphase. The MG132-treated cells were fixed, then subjected to immunostaining with anti-α-tubulin, anti-AcK250, and CREST (anti-ACA) antibodies. CREST and α-tubulin IF signals were analyzed using structured illumination microscopy (SIM) to determine the microtubule attachment (Fig. 1C). In SIM super-resolution imaging, unattached and attached kinetochores were assessed for the intensity and pattern of AcK250-BubR1 (Fig. 1C & Supplementary Fig. S1C). The immunofluorescence intensity of AcK250-BubR1 was the highest when both sister kinetochores were unattached. These AcK250-BubR1 displayed mostly crescent- and sometimes ring-shaped (Fig. 1C & D, No attachment). Chromosomes with a single attachment showed lower levels of AcK250-BubR1 (Fig. 1C & D, Single-attachment); AcK250-BubR1 was absent in amphitelically attached chromosomes (Fig. 1C & D, Bi-attachment).

Next, the IFA patterns of AcK250-BubR1 at the kinetochores were assessed after treating cells with various drugs perturbing KT-MT interactions: Nocodazole that disrupts microtubule polymerization; paclitaxel (Taxol) that interferes with depolymerization; Eg5 inhibitor Monastrol that induces tension-less KT-MT interactions; MG132 proteasome inhibitor that interferes with metaphase-anaphase transition due to inhibition of proteosome. The IFA intensity of AcK250 at kinetochores post-treatment was the highest in nocodazole-treated cells, followed by taxol>Monastrol>MG132 (Fig. 1E and F). Meanwhile, anti-BubR1 IFA levels in taxol, Monastrol, and MG132 treatments were not significantly different from each other. BubR1 levels at the kinetochore were the highest upon Nocodazole treatment (Fig. 1G and H).

### The volume of AcK250-BubR1 increases at the unattached kinetochore

The kinetochore is highly dynamic in size and shape, expanding through the formation of a fibrous meshwork known as the fibrous corona (38), which facilitates microtubule capture and chromosome biorientation (39). As K250 acetylation of BubR1 was associated with unattachment and increased volume of immunofluorescent signals (Fig. 1C), we investigated the relationship between BubR1 acetylation and kinetochore expansion.

First, the volume changes of BubR1 in relation to different KT-MT attachment statuses was assessed. Compared to kinetochores with no attachment (Fig. 2A & B, Supplementary Fig. S1C, No), the volume of BubR1 at kinetochore decreased by approximately 80% in monotelic attachments (Fig. 2A and B, Supplementary Fig. S1C, Single), and by approximately 63% in bipolarly attached kinetochores (Fig. 2A and B, Supplementary Fig. S1C, Bi). These results confirm that the previous report that BubR1 expands in unattached kinetochores (21).

**Figure 2.**
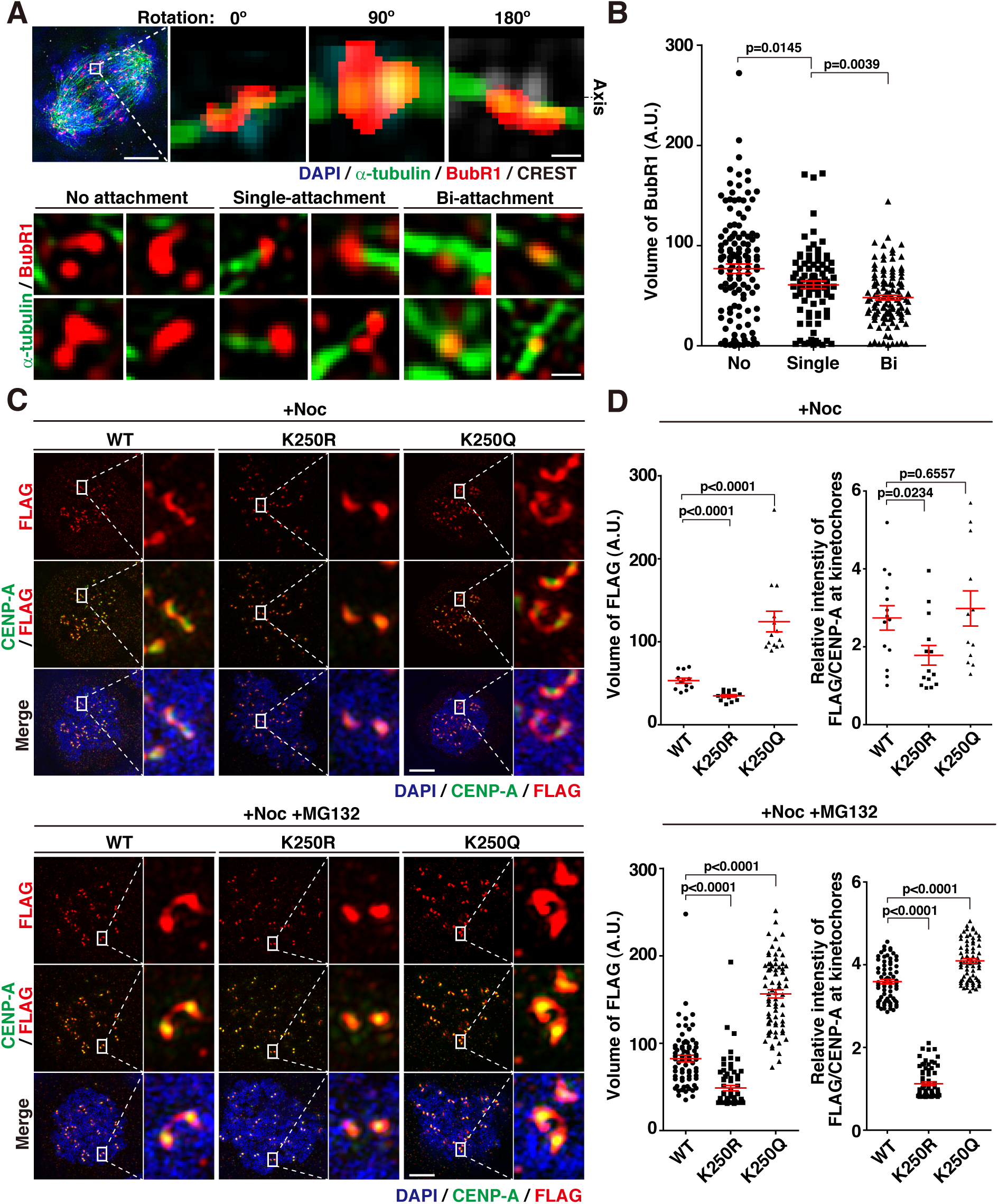
Lysine 250 acetylation of BubR1 (BubR1-AcK250) is associated with kinetochore expansion. (A) Representative images of M-phase HeLa cells immunostained with anti-BubR1, CREST, and α-tubulin antibodies. Cells were treated with 10 μM MG132 to enrich the metaphase population. Kinetochores were categorized into three groups based on 3D-reconstructed SIM images: No attachment, Single-attachment, or Bi-attachment. Scale bar, 5 μm. Bottom, 3D reconstruction. Scale bar, 0.5 μm. (B) Volume of BubR1 in cells exhibiting no attachment (No), monotelic attachment (Single), and amphitelic attachment (Bi). Number of chromosomes scored: No, n = 131; Single, n = 89; Bi, n = 115. Results are from three independent experiments (mean ± s.e.m.). (C) Representative SIM images of FLAG-tagged BubR1-WT, -K250R, and -K250Q in mitotic kinetochores. HeLa-FRT-TO cells expressing a single copy of FLAG-BubR1, -K250R, or -K250Q were depleted of endogenous BubR1 via siRNA transfection(34). Simultaneously, doxycycline was used to induce the expression of BubR1-WT, -K250R, and -K250Q, respectively. Cells were treated with nocodazole (Noc) alone or with MG132 (Noc + MG132) to assess the effect without interference from the stability of K250R. Cells were treated with nocodazole (200 ng/mL, 20 h) alone or in combination with MG132 (10 μM, final 2 h). Scale bar, 5 μm. (D) Volume of FLAG (Left) and relative intensities of FLAG to CENP-A (Right) in nocodazole-treated cells with or without MG132, as shown in the graphs. For nocodazole-only treatment: WT, n = 12; K250R, n = 14; K250Q, n = 14. For nocodazole + MG132 treatment: WT, n = 72, K250R, n = 72, K250Q, n = 72. The graph represents two independent experiments (mean ± s.e.m.).

Then we asked how acetylation of BubR1 influences kinetochore expansion. For this, we utilized HeLa-FRT-TO cells integrated with a single copy of FLAG-tagged wild-type BubR1 (*WT-BubR1*), acetylation-mimetic BubR1 (*K250Q*), or acetylation-deficient BubR1 (*K250R*) expression constructs (15,34). Expression of *WT-BubR1*, -*K250R*, and -*K250Q*, respectively was induced with doxycycline treatment. Endogenous BubR1 was depleted using siRNA targeting the 3’-UTR (Supplementary Fig. S2A and B). Forty-eight hours post-induction and simultaneous depletion of endogenous BubR1, cells were subjected to co-immunofluorescence with anti-FLAG (M2) and anti-CENP-A antibodies and examined by SIM super-resolution imaging (Fig. 2C). The volumes of the various BubR1 at the kinetochores were then measured.

The SIM analysis revealed that BubR1-K250 acetylation is associated with kinetochore expansion at the unattached kinetochores upon Nocodazole treatment: the IF intensity of acetylation-deficient BubR1 (K250R) was the lowest, while IF intensities of WT-BubR1 and the acetylation mimic K250Q were markedly higher (Fig. 2C & D, FLAG, relative intensity). The volumes of BubR1 variants were then measured. It showed that the volume of K250R was significantly smaller compared to that of wild-type, while the acetylation-mimic form (K250Q) exhibited a substantially larger volume (Fig. 2C and D). The difference in volume was more profound than the difference in intensity between K250Q and WT-BubR1 (Fig. 2C and D).

Acetylation/deacetylation of BubR1 at K250 functions as a molecular switch, converting BubR1 from an inhibitor to a substrate of APC/C-Cdc20 (11). Failure of BubR1-K250 acetylation results in premature degradation of BubR1 and consequent chromosome mis-segregation (15). We therefore assessed whether the decreased intensity and volume of K250R at unattached kinetochores was due to premature degradation of K250R-BubR1. To address this, the proteasome inhibitor MG132 was added during the final two hours of nocodazole treatment. Even with inhibition of protein degradation by co-treatment of MG132, the intensity and volume of K250R remained markedly lower compared to WT and K250Q (Fig. 2C and D, Noc+MG132). Intriguingly, the volume of K250Q was substantially higher compared to WT-BubR1, suggesting the possibility that K250 acetylation may directly promote kinetochore expansion.

### Acetylation of K250-BubR1 is required for CENP-E recruitment and fibrous corona expansion at the unattached kinetochore

Accurate chromosome segregation depends on the forces generated by end-on bipolar KT-MT interactions. In mitosis, chromosomes and plus- and minus-end motor proteins act in concert to achieve bipolar KT-MT attachment. End-on attachment is not achieved immediately; instead, chromosomes initially associate with the sides of existing kinetochore-fibers (K-fiber) and move to the equator before switching to end-on attachments. Polymerization of fibrous corona, the kinetochore expansion, facilitates the switch from lateral attachment to end-on attach (18,40,41).

Kinetochore expansion is driven by polymerization of the RZZ complex (Rod-ZW10-Zwilch), which appears in a crescent- or ring-shaped structure that surrounds the kinetochore. The polymerized RZZ complex recruits Mad1 and Mad2 (42). Upon successful end-on KT-MT attachment, the fibrous corona detaches. Since we observed that K250 acetylation was responsible for the crescent-shaped expansion of BubR1 in unattached kinetochores, we asked whether MAD2 and ZW10 expansion (17,19,21,22) was affected by BubR1 acetylation.

In SIM analysis of nocodazole-treated kinetochores, K250-BubR1 acetylation associated with crescent-shaped expansion of Mad2 and ZW10: Mad2 and ZW10 exhibited crescent-shaped expansion in cells expressing *WT*- and *K250Q-BubR1*, but not in *K250R*-expressing cells (Fig. 3A, +Noc). The crescent-shaped expansion was independent of the short half-life of K250R, as addition of MG132 did not alter the result (Fig. 3A, +Noc+MG132). Consistent with the formation of crescent-shaped architecture, the volumes of Mad2 and ZW10 were substantially higher in *K250Q*-expressing cells compared to *K250R*-expressing cells (Supplementary Fig. S3A).

**Figure 3.**
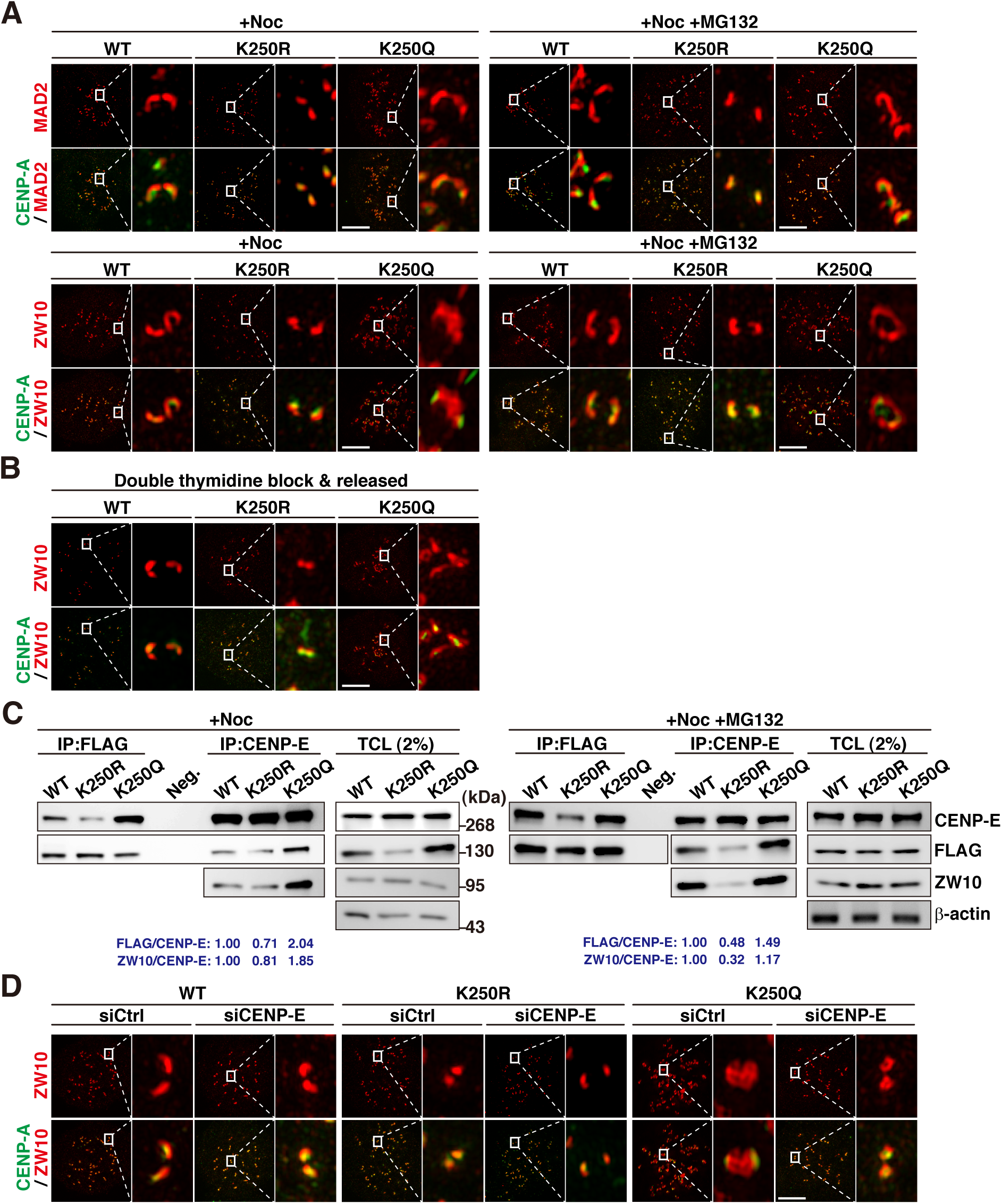
K250-BubR1 acetylation is involved in the maintenance of crescent-shape fibrous corona. (A) Representative images of immunostaining with anti-Mad2 and anti-ZW10 antibodies after induction of BubR1-WT, -K250R, and -K250Q expression in HeLa-FRT-TO cells. Nocodazole (200 ng/mL) alone or with MG132 was applied before fixation. Endogenous BubR1 was depleted via siRNA. Number of cells scored for Mad2: WT, n = 15; K250R, n = 20; K250Q, n = 18. For ZW10: WT, n = 12; K250R, n = 14; K250Q, n = 14 (mean ± s.e.m.). Scale bar, 5 μm. (B) Representative SIM images of immunostained ZW10 at kinetochores of HeLa cells expressing FLAG-tagged BubR1-WT, -K250R, or -K250Q following double thymidine block and release. FRT/TO HeLa cells were treated with doxycycline for 55 hours to induce expression of ectopic BubR1 variants. Endogenous BubR1 was depleted using siRNA targeting the 3’UTR. Cells were fixed for immunostaining 10 h after the final thymidine release to enrich the mitotic population. (C) Effects of BubR1 acetylation status on CENP-E and RZZ complex formation. Cells were treated with nocodazole alone (+Noc) or with MG132 (+Noc +MG132) to assess the binding affinity without interference from the stability of K250R. Relative protein intensities were measured using a densitometer and are shown in blue. (D) Representative images showing immunostained ZW10 and anti-CENP-A antibodies in FRT/TO HeLa cells expressing BubR1-WT, K250R, or K250Q with (siCtrl) or without CENP-E (siCENP-E). Cells were transfected with siRNA targeting luciferase (siCtrl) or CENP-E (siCENP-E) for 48 hours prior to fixation. Nocodazole was administered for 20 hours before analysis.

To assess the fibrous corona during a normal cell cycle, minimizing the effects from microtubule poison, cells were synchronized in S phase by double thymidine block then washed to release into mitosis (Supplementary Fig. S3B). When prometaphase cells were subjected to SIM analysis, it confirmed that K250-BubR1 acetylation deficiency interferes with fibrous corona polymerization (Fig. 3B). Treatment of cells with the pan-HDAC inhibitor Trichostatin A (TSA) and nocodazole also showed that Mad2 and ZW10, as well as AcK250-BubR1, increase their volume around the kinetochore in a crescent shape, supporting that maintaining AcK250-BubR1 is interlinked with kinetochore expansion (Supplementary Fig. S3C).

BubR1 associates with CENP-E motor protein (43,44), which facilitates chromosome congression by mediating the lateral sliding of chromatids along the existing K-fibers (45). We previously showed that K250 acetylation affects the binding to CENP-E (15). As K250 acetylation was associated with ZW10 and Mad2 expansion at the outer kinetochore, we asked whether the association of BubR1 with CENP-E, which depends on acetylation status, affected ZW10 volume expansion. Immunoprecipitation followed by Western blotting confirmed that the binding of acetylation-deficient BubR1 (K250R) to CENP-E decreased to 0.71, while the binding of acetylation-mimetic K250Q increased to 2-fold in nocodazole-treated cells, compared to WT (Fig. 3C, +Noc). In the same lysate, IP with anti-CENP-E and WB with anti-ZW10 antibodies revealed that ZW10 interaction with the CENP-E was reduced to 0.8-fold in K250R-expressing cells, whereas K250Q-expressing cells showed ∼1.9-fold increase in CENP-E and ZW10 interaction (Fig. 3C, +Noc). The decreased CENP-E association with ZW10 in *K250R*-expressing cells was minimally affected by premature degradation of K250R, as the addition of MG132 led to similar results (Fig. 3C, +Noc+MG132).

In addition, depletion of CENP-E abolished the crescent shape as well as decreased volume of ZW10 in *WT-BubR1* and *K250Q*-expressing cells to the levels comparable to those observed in *K250R*-expressing cells with CENP-E (Fig. 3D and Supplementary Fig. S3D). These results indicate that acetylation of K250 of BubR1 links fibrous corona polymerization and CENP-E recruitment to the outer kinetochore.

### Phosphorylation by Aurora B kinase at the TPR domain precedes K250 acetylation of BubR1

We next sought to reveal how the signal of unattachment is transduced to K250-BubR1 acetylation. For this, we hypothesized that a certain mitotic kinase may be involved in BubR1 acetylation. The AurkB plays a pivotal role in proper KT-MT attachment. It is known that at the inner kinetochore, AurkB phosphorylates Ndc80 of the KMN network (28,46) and releases incorrect attachment (47). When proper end-on attachment is achieved, phosphatases PP1 and PP2A antagonize excessive AurkB activity and stabilize KT-MT attachment for anaphase onset (48,49).

Plk1 phosphorylates BubR1 in a tension-dependent manner (50). Phosphorylation by Plk1 at the KARD domain at the C-terminus of BubR1 recruits PP2A-B56α, which antagonizes excessive Aurora B kinase activity at the kinetochore, thereby stabilizing microtubule attachments (37). It should be noted that K250 acetylation, located distally from the KARD domain, is crucial for the stabilization of KT-MT attachments (15).

The MPS1 kinase is required for the establishment of fibrous corona by phosphorylating the dynein adaptor Spindly, leading to conformational change of spindle and subsequent oligomerization of RZZ complex (39). MPS1 also recruits Mad1 and Mad2 to kinetochores, establishing the initiation of SAC activity (51,52).

To assess which mitotic kinases and how they signal to BubR1 acetylation, HeLa cells were treated with various mitotic kinase inhibitors at the final two hours (Supplementary Fig. S4A) of nocodazole treatment and subjected to IFA with anti-AcK250 mAb. The volume expansion and relative intensity of AcK250 was inhibited by AurkB inhibitor, Hesperadin or ZM447439, but not with the Plk or MPS1 inhibitor, BI2536 and Reversine (Fig. 4A and B). Previous reports showed that BubR1 expansion is enhanced by the phosphatase 1 inhibitor I-2 (21), and that AurkB pathway controls the conversion of lateral to end-on attachment via BubR1 recruiting PP2A (53). These results suggested that AurkB may be involved in K250-BubR1 acetylation.

**Figure 4.**
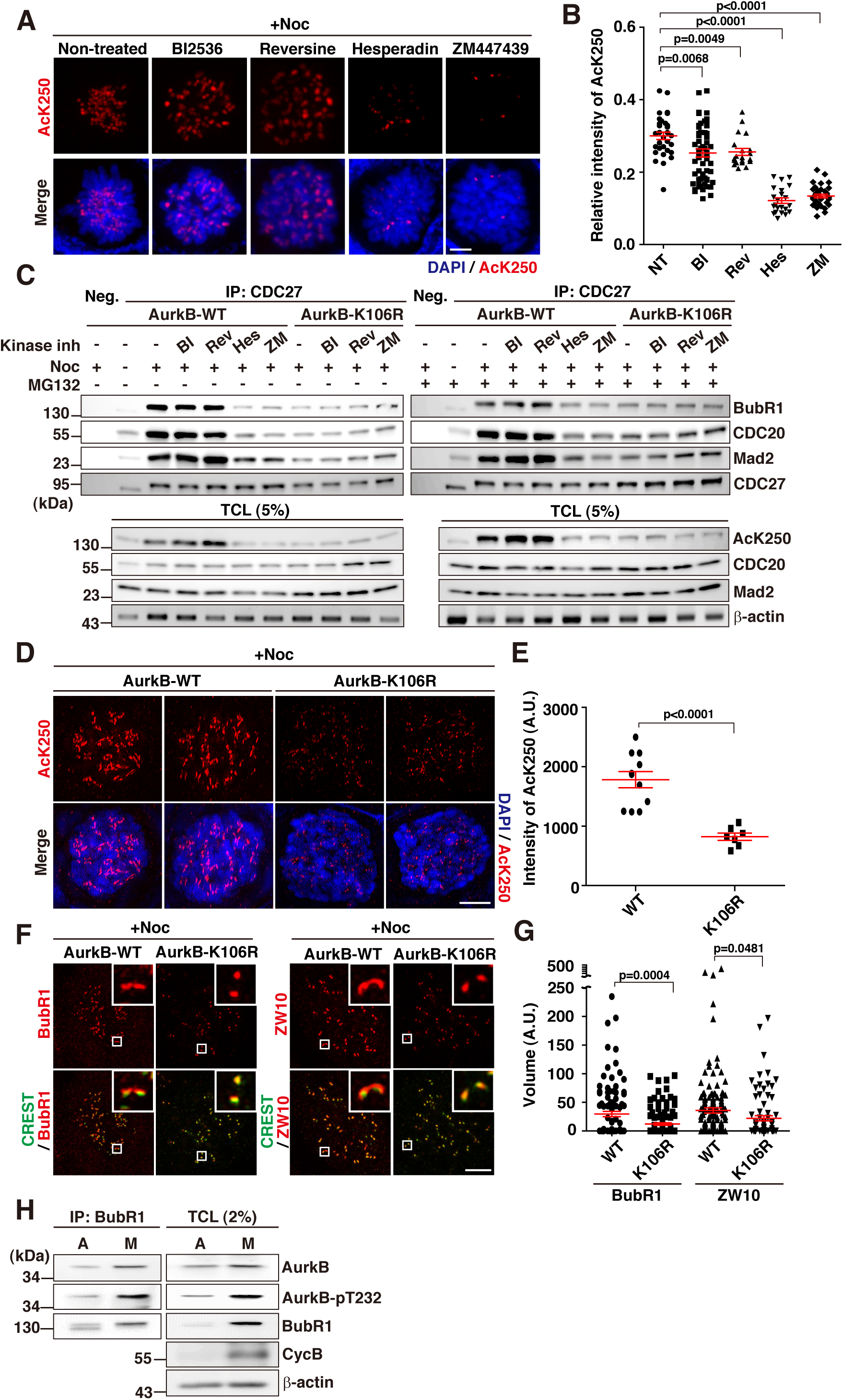
Aurora kinase B activity is required for BubR1 acetylation and maintenance of fibrous corona. A) Immunostaining with anti-AcK250 mAb after treatment of nocodazole-arrested cells with the indicated mitotic kinase inhibitors: 10 nM BI2536 (Plk1 inhibitor), 2 μM Reversine (Mps1 inhibitor), 200 nM Hesperadin (Aurora B inhibitor), and 2 μM ZM447439 (Aurora B inhibitor). The kinase inhibitors were applied during the final 2 hours of a 20-hour nocodazole treatment. Scale bar, 5 μm. (B) Relative intensities of AcK250 at kinetochores from (A): Nocodazole-arrested cells without kinase inhibitor addition (NT), and nocodazole + BI2536-treated (BI2536), Reversine-treated (Rev), Hesperadin-treated (Hes), or ZM447439-treated (ZM) cells (mean ± s.e.m.). Results are from two independent experiments. (C) Assessing the effect of Aurora B kinase activity on MCC maintenance. HeLa cells were transfected with Aurora B-WT or kinase-dead mutant K106R-expressing constructs. Endogenous Aurora B was depleted using siRNA targeting the 3’UTR. BI2536 (10 nM), Reversine (2 μM), Hesperadin (200 nM), or ZM447439 (2 μM) were added during the final 2 hours of a 20-hour nocodazole treatment (Left). Immunoprecipitation was performed using an anti-CDC27 (APC3) antibody, followed by WB with indicated antibodies. To assess MCC stability, MG132 was added with the indicated kinase inhibitors in nocodazole-arrested cells (Right). WB of the total cell lysate (5%) is shown for the loading control (Bottom). (D) Representative images of anti-AcK250 immunostaining in cells expressing wild-type Aurora B (AurkB-WT) or kinase-dead Aurora B mutant (AurkB-K106R). Scale bar, 5 μm. (E) Intensity of AcK250 measured from (D). Number of cells counted: WT, n = 11; AurkB-K106R, n = 7 (mean ± s.e.m.). (F) Effect of Aurora B kinase activity on the expansion of BubR1 (Left) and ZW10 (Right) at nocodazole-treated kinetochores. Scale bar, 5 μm. (G) Volume of BubR1 and ZW10 at the kinetochores from (F). Number of cells scored: BubR1 in WT, n = 105; BubR1 in K106R, n = 121; ZW10 in WT, n = 153; ZW10 in K106R, n = 85 (mean ± s.e.m.). (H) Interaction between BubR1, Aurora B, and phosphorylated Aurora B (AurkB-pT232) was assessed by IP with anti-BubR1 and WB with indicated antibodies. Mitotic HeLa cells were collected by mitotic shake-off after 20 h of nocodazole treatment (M). Interphase attached cells (A) were used as a control.

To corroborate whether AurkB kinase activity impinges on BubR1 acetylation, MCC maintenance was analyzed. Previously, we have shown that BubR1 acetylation is required for MCC maintenance but not in initial MCC assembly (15). Immunoprecipitation with anti-CDC27, a component of APC/C, followed by Western blotting showed that MCC maintenance required AurkB activity, as Hesperadin and ZM447439 treatment abolished MCC bound to CDC27. PLK1 or MPS1 inhibitors did not affect MCC maintenance, as the kinase inhibitors were applied in the last two hours of nocodazole treatment (Fig. 4C, AurkB-WT). The expression of a kinase-dead mutant of AurkB (AurkB-K106R) (54–57) abolished MCC even under nocodazole treatment. Addition of MG132 led to similar results (Fig. 4C, +Noc+MG132). Note that western analysis of total lysates confirmed that acetylation of K250 in BubR1 was abolished upon AurkB inhibition, both by AurkB inhibitor treatment and kinase dead mutant AurkB expression (Fig. 4C, TCL, AcK250).

We then asked whether AurkB activity is involved in the crescent-shaped kinetochore expansion of AcK250-BubR1. HeLa cells were depleted of endogenous Aurora B (Supplementary Fig. S4B) and were transfected with *AurkB* (*AurkB-WT*)-or a kinase-dead (*AurkB-K106R*)-expression constructs (54–57). Expression of *AurkB-K106R* markedly reduced the intensity and volume of AcK250-BubR1 compared to *AurkB-WT*-expressing prometaphase cells (Fig. 4D and E). Moreover, *AurkB-K106R* expression led to a marked decrease in the volume of ZW10, as well as BubR1 (Fig. 4F and G, see insets). These results suggest that AurkB activity is required for K250-BubR1 acetylation and the subsequent crescent-shaped expansion of BubR1 and the fibrous corona. Importantly, the effect of *AurkB-K106R* expression on the reduction of AcK250-BubR1 was more profound than its effect on total BubR1 (Figure S4C).

In binding assays utilizing IP and WB, BubR1 interacted with the kinase active form of AurkB (AurkB-pT232) (58) specifically in mitosis but markedly less in non-mitotic cells (Fig. 4H). In *in vitro* analysis, the recombinant AurkB was capable of phosphorylating *in vitro*-translated BubR1 (Supplementary Fig. S4D). These results demonstrate that AurkB-mediated phosphorylation is required for K250-BubR1 acetylation in mitosis. The AurkB-mediated phosphorylation then acetylation of BubR1 is associated with the polymerization of fibrous corona in unattached kinetochore.

### AurkB phosphorylates S16 and S39 in BubR1, enabling K250 acetylation

Next, we aimed to identify the phosphorylation sites in BubR1 targeted by AurkB. To this end, HeLa-FRT-TO cells expressing FLAG-tagged wild-type BubR1 (*WT*), *K250R*, and *K250Q*, respectively, were treated with nocodazole, immunoprecipitated with anti-FLAG monoclonal antibody, then subjected to in-gel digestion for mass spectrometry analysis. The liquid chromatography-tandem mass spectrometry (LC-MS/MS) identified seventy-nine phosphorylation sites in BubR1. After repeating the LC-MS/MS three times, fifteen candidate sites were selected for further analysis (Fig. 5A and Supplementary Table S1). The residues identified in BubR1-WT and BubR1-K250Q, but not in K250R, were prioritized (Supplementary Fig. S5A). Known Plk-1 phosphorylation sites were excluded, as K250-acetylation was not affected by Plk1 inhibition (Fig. 4A-C).

**Figure 5.**
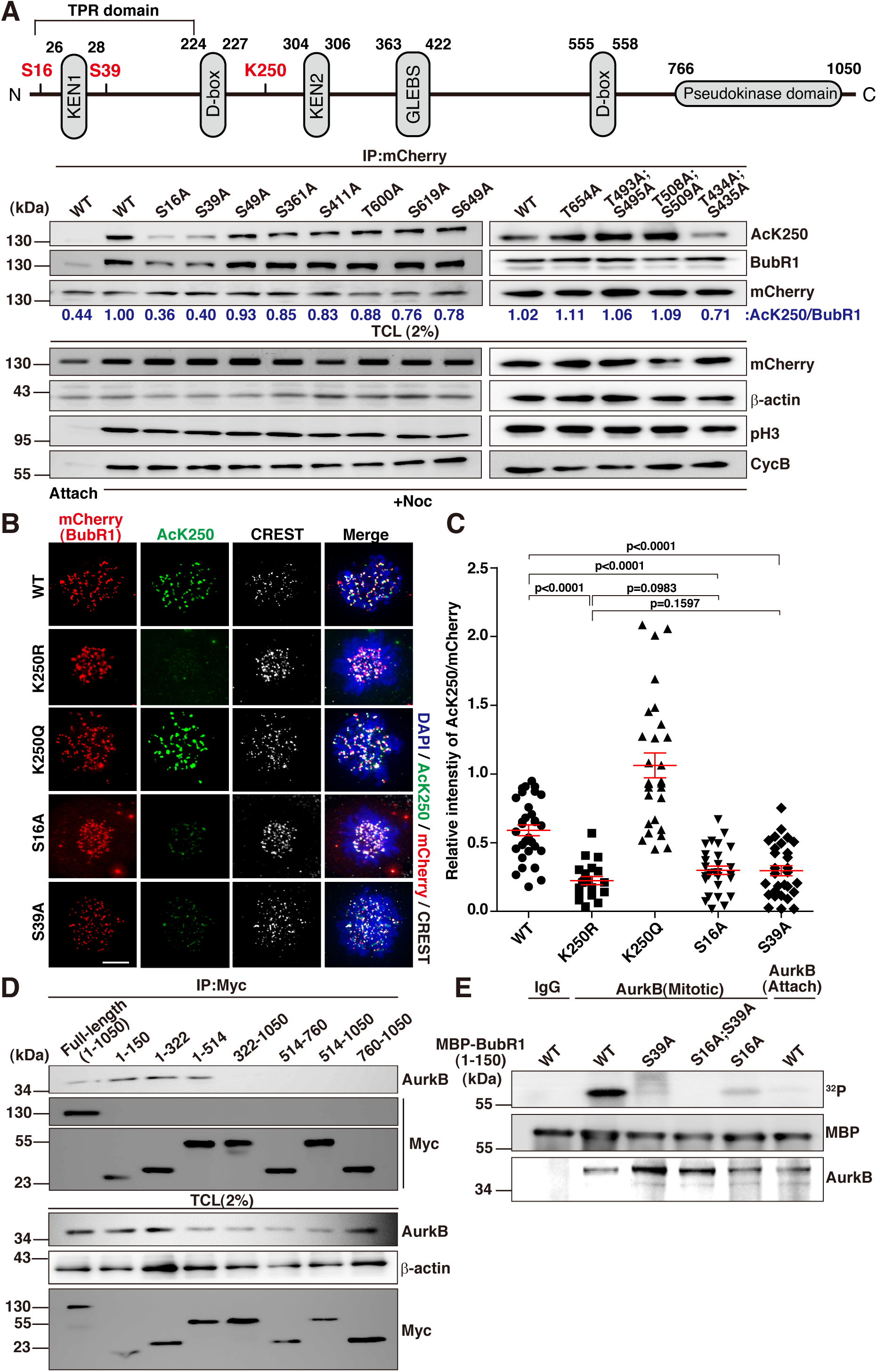
Aurora B phosphorylates Serine 16 and Serine 39 of BubR1 upon nocodazole treatment, which is required for K250 acetylation. (A) Identification of Aurora B-mediated phosphorylation sites of BubR1 in response to nocodazole treatment. (Top) Schematic illustration of BubR1, indicating potential Aurora B-mediated phosphorylation sites (Ser16 & Ser39) and the K250 acetylation site. (Bottom) IP-WB analysis to identify phosphorylation sites that crosstalk with K250 acetylation. mCherry-tagged BubR1 expression constructs were mutagenized in vitro to substitute the phosphorylation sites with alanine, based on proteome analysis (Supplementary Fig. S5), and then transfected into HeLa cells. Cells were treated with nocodazole (200 ng/mL) for 20 h and mitotic cells were collected by shake-off. Attached cells (Attach) were used as a control. Mitotic cell lysates, along with control lysates from attached cells (WT, Attach), were subjected to immunoprecipitation with anti-mCherry antibody, followed by WB with anti-AcK250 mAb. The same blot was reprobed with anti-BubR1 and anti-mCherry antibodies for normalization. Total cell lysates were subjected to WB with anti-phospho-H3 and anti-cyclin B antibodies to assess whether the cells were in mitosis. WB with anti-mCherry and anti-β-actin antibodies in TCL were used as a loading control. Relative band intensities (AcK250/BubR1) were measured using a densitometer and are indicated. (B) Effect of Ser16 or Ser39 phosphorylation on K250 acetylation. Immunofluorescence assay in cells expressing the indicated mCherry-tagged BubR1 variants. Nocodazole-treated cells were formaldehyde-fixed and subjected to immunostaining with anti-AcK250, anti-ACA (CREST), and anti-mCherry antibodies. (C) Graph showing the intensities of anti-AcK250 immunofluorescence from (B). Anti-AcK250 immunofluorescence signals were normalized to anti-mCherry. The results are from two independent experiments (WT, n = 30; K250R, n = 20; K250Q, n = 24; S16A, n = 28; S39A, n = 29) (mean ± s.e.m.). (D) Identification of the Aurora B-binding region in BubR1. IP with 9E10 (anti-Myc) and WB with anti-AurkB (Aurora B kinase) were performed. All BubR1-expressing constructs were Myc-tagged. The same blot was reprobed with 9E10 for normalization. Two percent of TCL was subjected to WB with the indicated antibodies. (E) In vitro phosphorylation assay to confirm the phosphorylation of S16 and S39 by AurKB. Recombinant BubR1 (1-150 a.a.), purified from E. coli, was subjected to in vitro phosphorylation by Aurora B immunoprecipitated from nocodazole-arrested HeLa cells. Attached cells were used as a control (Attach). [γ-32P]-ATP was used in the assay to facilitate detection. WB with anti-MBP and anti-AurkB show the loading control.

The candidate phosphorylation sites were then mutated to alanine using *in vitro* mutagenesis, and the resultant constructs tagged with mCherry were transfected into HeLa cells. Forty-eight hours post-transfection, cells were treated with nocodazole for the final twenty hours. Then the cells were subjected to immunoprecipitation with anti-mCherry antibody, followed by Western analysis with anti-AcK250-BubR1 or anti-BubR1 antibodies (Fig. 5A).

Through these serial experiments, serines 16 and 39 were identified as phosphorylation sites associated with K250 acetylation, as the S16A and S39A mutants failed to undergo K250-BubR1 acetylation in mitotic cells (Fig. 5A). In mass spectrometry, these sites were phosphorylated in WT-BubR1 and K250Q, but not in K250R-expressing cells upon nocodazole treatment (Supplementary Fig. S5B). Located within the tetratricopeptide repeat (TPR) domain of BubR1(59), which interacts with KNL1 (Blinkin) (60,61), S16 and S39 are positioned adjacent to the KEN1 degron (Fig. 5A, top).

The serine 39 of BubR1 conforms to the known consensus sequence for AurkB substrates, which typically includes basic residues two or three positions upstream of the phosphorylation site (56,62–64). Meanwhile, S16 (SEAMS16L17) does not adhere to the canonical motif. However, some Aurora B substrates such as MgcRacGAP (65), Dsn1 (30,66), and INCENP (56,64) lack this motif but are nevertheless phosphorylated by AurkB, suggesting that S16 might also be phosphorylated by AurkB along with the canonical S39 when unattached.

Next, the effects of S16 or S39 phosphorylation in BubR1 acetylation were assessed by IFA with anti-AcK250 mAb in prometaphase cells after transfection with mCherry-tagged *BubR1* variants. Both S16A and S39A mutants exhibited defects in K250 acetylation in nocodazole-treated prometaphase cells, comparable to the K250R mutant (Fig. 5B, AcK250). Cells expressing wild-type (*WT*) and acetylation-mimetic (*K250Q*) BubR1 displayed robust AcK250 immunofluorescence at the kinetochores (Fig. 5B and C). These data indicate that phosphorylation of S16 and/or S39 is interconnected with K250 acetylation.

In the *in vitro* analysis of AurkB-interacting domain in BubR1 by employing BubR1 deletion constructs tagged with Myc, the IP and WB analysis confirmed that the N-terminus of BubR1 (amino acids 1-150), which contains S16 and S39, indeed interacted with Aurora B (Fig. 5D).

To confirm S16/S39 are indeed phosphorylated by AurkB upon unattachment, recombinant MBP-tagged BubR1 (amino acids 1-150, Supplementary Fig. S5C) variants were subjected to an *in vitro* phosphorylation assay with AurkB immune-complex obtained from nocodazole-treated HeLa cells (Fig. 5E). The [γ-32P]-ATP-labeled *in vitro* phosphorylation assay showed that WT-BubR1 (1-150) was phosphorylated by AurkB. In contrast, S16A and S39A mutants, as well as the S16A;S39A double mutant, were not phosphorylated by AurkB. The AurkB immunoprecipitates from interphase cells, obtained from the remaining attached cells after nocodazole treatment followed by mitotic shake-off (Fig. 5E, AurkB Attach), failed to phosphorylate BubR1 (1-150). These results confirm that AurkB phosphorylates BubR1 S16 and/or S39 in mitosis. Finally, *in vitro* phosphorylation assays employing *in vitro*-translated Myc-BubR1 variants and recombinant AurkB confirmed that S16 and S39 are indeed recognized by AurkB (Supplementary Fig. S5D). Taken together, these data support the conclusion that AurkB phosphorylates S16 and S39, which then promotes K250 acetylation of BubR1 in unattached kinetochores.

### AurkB-mediated phosphorylation then acetylation on BubR1 constitutes the SAC signaling

We next investigated the impact of S16 and S39 phosphorylation defects in MCC maintenance and mitotic timing. HeLa cells transfected with various mCherry-tagged BubR1 constructs were subjected to nocodazole treatment. Twenty hours post-treatment, cells underwent to IP with anti-mCherry antibody, followed by WB with anti-CDC20, -Mad2, and -Bub3 antibodies and assessed for MCC stability. Like K250R-BubR1 (11,34), the BubR1 S16A and S39A mutants failed to maintain MCC, while other mutants were unaffected (Fig. 6A, +Noc). Co-treatment with the proteasome inhibitor MG132 during the last two hours of nocodazole treatment restored MCC levels in cells expressing S16A and S39A, indicating that phosphorylation at these sites is related to BubR1 stability critical for MCC maintenance (Fig. 6A, +Noc+MG132).

**Figure 6.**
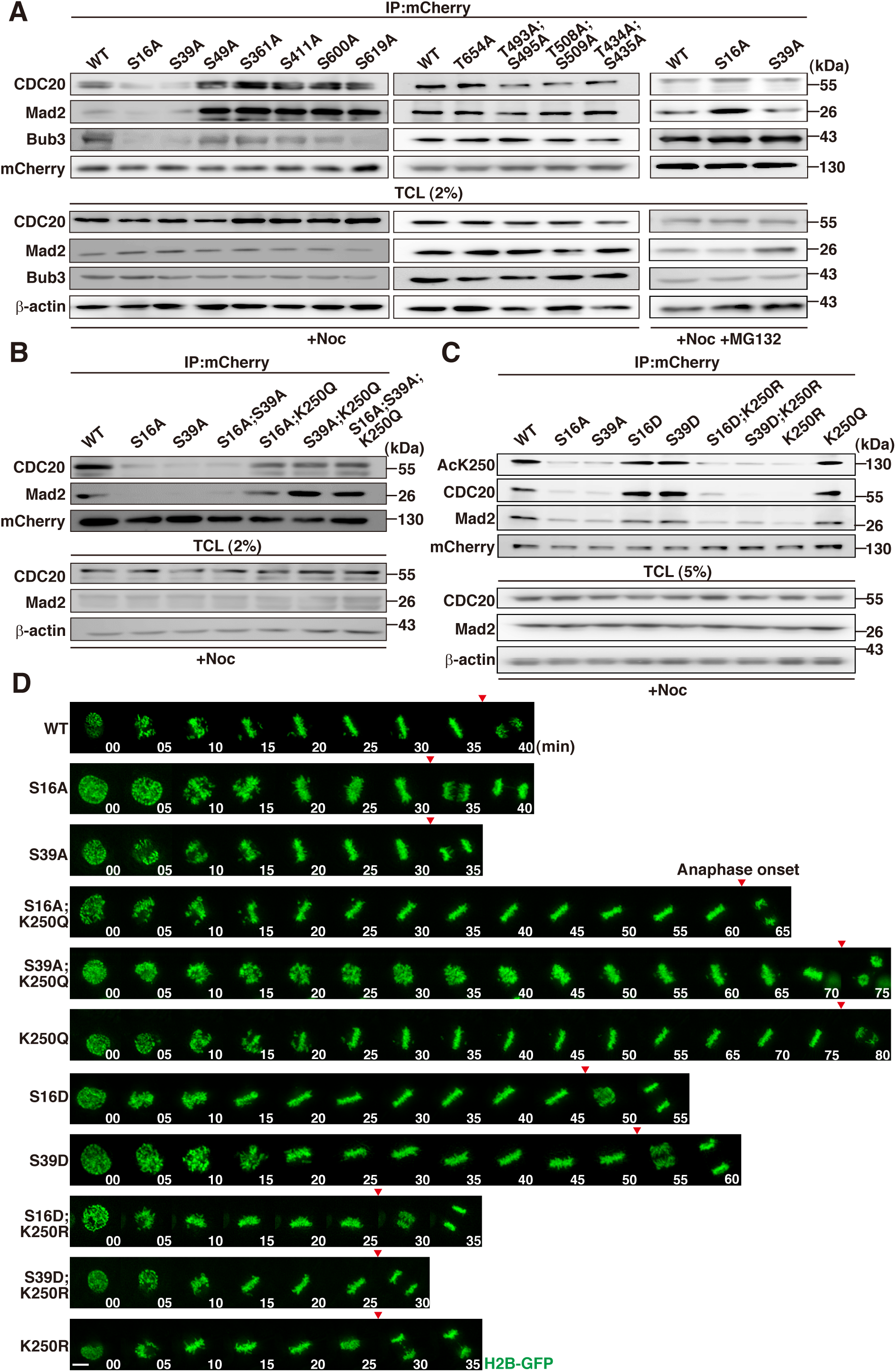
Serines 16 and 39 phosphorylation by AurKB is required for K250 acetylation in SAC signaling. (A) IP-WB analysis to assess the effect of Ser16 and Ser39 phosphorylation on MCC maintenance after nocodazole treatment. The indicated mCherry-tagged BubR1 variants were transfected, and endogenous BubR1 was depleted by co-transfection of siRNA for BubR1 targeting the 3’UTR. Immunoprecipitation was performed with an anti-mCherry antibody, followed by WB with the indicated antibodies. Results for nocodazole-arrested HeLa cells treated with MG132 for 2 hours before harvest are shown at right (+Noc +MG132). WB of 2% TCL is shown at the bottom. (B, C) Effects of phosphorylation and acetylation on MCC maintenance. (B) Effects of AurKB-mediated phosphorylation at Serines 16 and 39 on MCC maintenance. Phospho-deficient mutants S16A-BubR1 and S39A-BubR1, as well as the S16A;S39A mutant, and phospho-deficient mutations introduced into the acetylation-mimetic form (S16A;K250Q, S39A;K250Q, S16A;S39A;K250Q) were transfected into HeLa cells, along with siBubR1 to deplete endogenous BubR1. The cells were treated with nocodazole and subjected to IP with anti-mCherry antibody, followed by WB with the indicated antibodies. (C) Effect of the phospho-mimetic S16D-BubR1 or S39D-BubR1 on MCC maintenance. The acetylation-deficient K250R mutation was introduced into S16D or S39D (S16D;K250R and S39D;K250R) variants, and their capabilities for maintaining MCC after nocodazole treatment were assessed. All BubR1 variants were tagged with mCherry, transfected into HeLa cells, and subjected to IP and WB using the indicated antibodies. (D) Effect of AurKB-mediated phosphorylation and K250 acetylation on mitotic timing. Endogenous BubR1 was depleted using siRNA, and the indicated mCherry-tagged BubR1 variants were transfected into HeLa cells stably expressing H2B-GFP (11). Images were captured at 5-minute intervals. Arrows mark anaphase onset.

Then we asked whether S16 and/or S39 phosphorylation precedes K250 acetylation. For this, we introduced the *K250Q* mutation into the *S16A* and *S39A* mutants and then assessed for MCC stability. The results showed that MCC maintenance upon nocodazole treatment, measured by the levels of CDC20 and Mad2 complexed with BubR1 variants, restored to the levels comparable to WT when the acetylation-mimetic mutation was introduced to phospho-defective mutants (S16A;K250Q, S39A;K250Q, S16A;S39A;K250Q), while S16A, S39A, or S16A; S39A mutants failed to maintain MCC (Fig. 6B, left).

When phospho-mimetic BubR1 mutations (S16D, S39D) were introduced into *K250R*, the resulting proteins failed to sustain MCC levels, whereas the S16D and S39D mutants maintained MCC (Fig. 6C). These findings indicate that S16 and S39 phosphorylation by AurkB is a prerequisite for K250 acetylation. Subsequent K250-BubR1 acetylation stabilizes MCC.

The effect of phosphorylation and acetylation in mitotic timing and chromosome segregation were then assessed. Compared to WT, mitotic timing, as measured from nuclear envelope breakdown (NEBD) to anaphase onset, was shortened by approximately 5 minutes in cells expressing *S16A* and *S39A* mutants (Fig. 6D, S16A, S39A and Supplementary Video S1-S3). When the *K250Q* mutation was introduced into these phospho-defective mutants, mitotic timing was elongated by approximately 25-35 minutes (Fig. 6D, S16A;K250Q, S39A; K250Q and Supplementary Video S4-S5), similar to *K250Q*-mutant-expressing cells (Fig. 6D, K250Q and Supplementary Video S6).

When S16D and S39D mutants were expressed, mitotic timing was elongated by 15-20 minutes compared to WT (Fig. 6D, S16D, S39D and Supplementary Video S7-S8). Introduction of *K250R* into the S16D and S39D mutants substantially shortened mitotic timing by approximately 5-10 minutes compared to WT (Fig. 6D, S16D;K250R or S39D;K250R and Supplementary Video S9-S10), comparable to the timing observed in *K250R* expressing cells (Fig. 6D, K250R and Supplementary Video S11). Endogenous BubR1 was depleted in all experiments. We observed mis-segregation of chromosomes, including lagging chromosomes and segregation without congression, in cells expressing *S16A*, *S39A*, *K250R*, *S16D*; *K250R*, and *S39D;K250R*. Conversely, introduction of *K250Q* into the S16A and S39A mutants eliminated chromosome mis-segregation, suggesting that S16 and S39 phosphorylation is a prerequisite for acetylation of K250-BubR1; K250 acetylation is critical in SAC activity and deacetylation of K250 triggers the onset of anaphase.

### Stepwise S16/S39 phosphorylation then K250 acetylation in BubR1 aligns with kinetochore expansion

We next investigated whether ser16 and ser39 phosphorylation of BubR1 by AurkB affects kinetochore expansion. Structured illumination microscopy (SIM) analysis revealed that volume expansion of Bub1 (67) and Mad2 (42,67) was impaired in cells deficient in S16 and S39 phosphorylation (Fig. 7A). Likewise, the crescent-shaped volume increase of ZW10 was disrupted in BubR1-S16A- and -S39A-expressing cells under nocodazole treatment (Fig. 7B and C). When the K250Q mutation was introduced into the S16A and S39A mutants, it restored the volume increase and crescent-shaped expansion (Fig. 7B and C, S16A; K250Q or S39A; K250Q). Consistent with the MCC maintenance and mitotic timing data, introduction of *K250R* into the *S16D* and *S39D* mutants resulted in the loss of volume and crescent-shaped expansion observed in *S16D*- and *S39D*-mutant-expressing nocodazole-treated cells (Fig. 7B and C). Collectively, these results indicate that Aurora B-mediated phosphorylation of S16 and S39 sets the stage for K250 acetylation, constituting a checkpoint signaling code that transduces unattachment signal to fibrous corona polymerization and MCC maintenance until bi-polar end-on attachment is accomplished.

**Figure 7.**
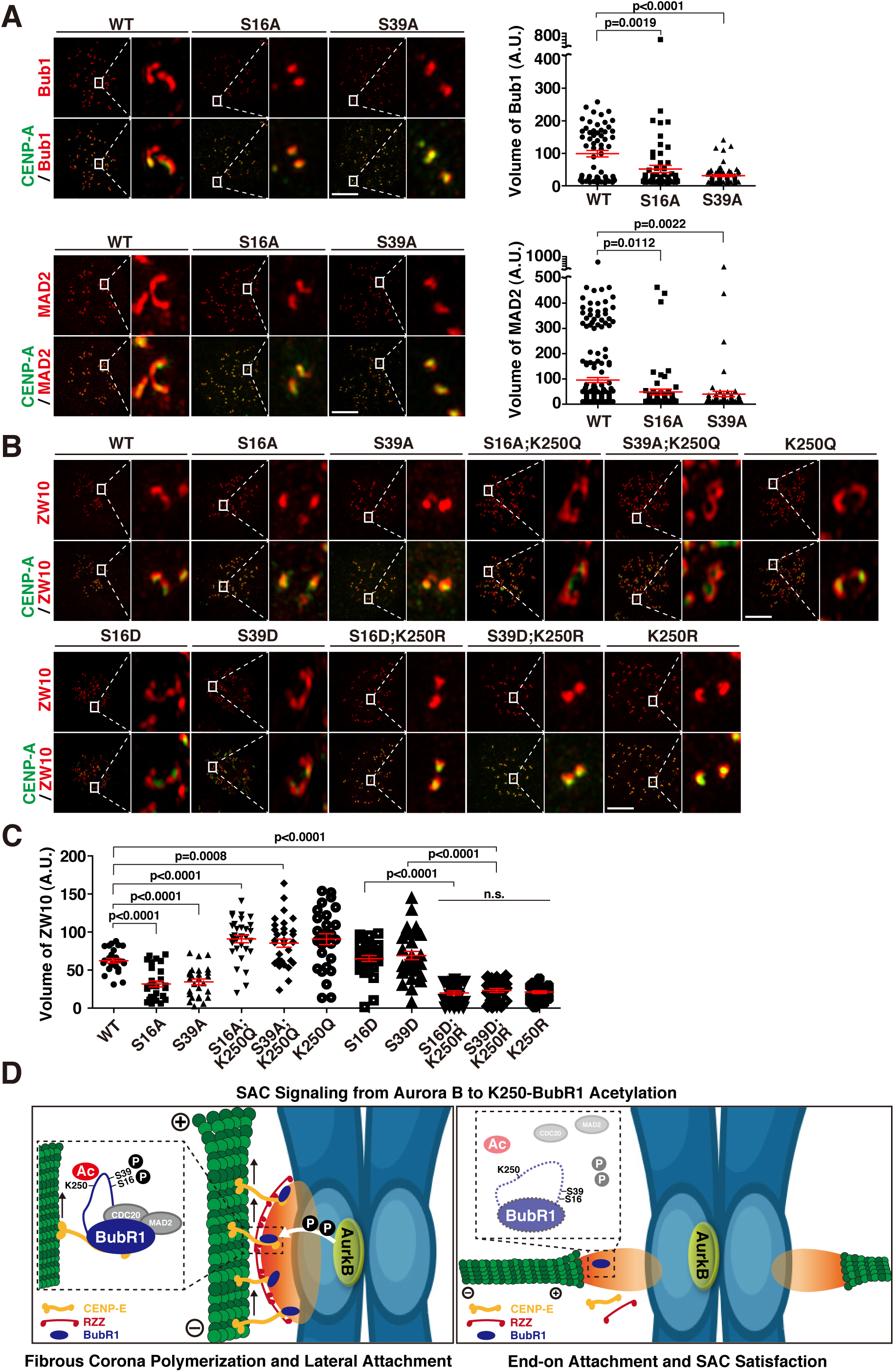
AurkB-AcK250-BubR1 pathway coordinates the unattachment signal to the maintenance of fibrous corona and SAC. (A, B) Effects of AurkB-mediated phosphorylation and subsequent K250 acetylation on fibrous corona expansion. (A) SIM images of anti-Bub1 or anti-MAD2 immunostaining in nocodazole-treated cells are shown. The indicated BubR1 constructs were transfected with siBubR1 targeting the 3’UTR 48 hours before fixation, and nocodazole (200 ng/mL) was applied for 20 hours. Volumes were measured from the immunofluorescence assays shown. Number of cells scored: Bub1 in WT, n = 63; Bub1 in S16A, n = 64; Bub1 in S39A, n = 70; MAD2 in WT, n = 196; MAD2 in S16A, n = 196; MAD2 in S39A, n = 69 (mean ± s.e.m.). (B) SIM images of anti-ZW10 immunostaining in nocodazole-treated cells. Endogenous BubR1 was depleted via co-transfection with siBubR1. Nocodazole (200 ng/mL) was administered for 20 hours after 48 h of transfection. Insets show enlarged images of ZW10 and CREST staining. Scale bar, 5 μm. (C) Volume of ZW10 measured from the immunofluorescence assay shown in (B). Number of cells scored: WT, n = 27; S16A, n = 29; S39A, n = 27; S16A;K250Q, n = 31; S39A;K250Q, n = 32; K250Q, n = 29; S16D, n = 31; S39D, n = 32; S16D;K250R, n = 27; S39D;K250R, n = 27; K250R, n = 27 (mean ± s.e.m.). (D) Model illustrating the coordination of AurKB-mediated phosphorylation and K250 acetylation in spindle checkpoint signaling. At unattached kinetochores, AurKB phosphorylates BubR1 at S16 and S39, which promotes BubR1 acetylation at K250 by PCAF. This acetylation ensures the maintenance of the RZZ complex and MCC. It also sustains CENP-E at the kinetochore, preparing for switch from lateral to end-on attachment (Left). Upon successful end-on capture, PP2A activity recruited by AckK250-BubR1 antagonizes AurkB activity, stabilizing KT-MT attachment. Simultaneously, BubR1 immediately deacetylates and degraded through APC/C-mediated ubiquitination, resulting MCC disassembly and SAC silencing.

## DISCUSSION

With compelling lines of evidence, we have demonstrated that AurkB-mediated phosphorylation at S16 and S39 of BubR1 enables K250 acetylation in an unattachment-dependent manner. This stepwise phosphorylation-acetylation checkpoint signaling supports the the fibrous corona polymerization with MCC maintenance, coordinating lateral to end-on attachment with SAC-regulated mitotic timing (Fig. 7D, left). When appropriate bipolar endon attachment is accomplished, BubR1 is deacetylated (34), leading to disentaglement of fibrous corona and disassembly of MCC due to degradation of deacetylated BubR1, which is also shown by K250R (Fig. 7D, Right).

When a kinetochore is unattached, the fibrous corona, RZZ-Spindly complex and Mad1, Mad2 included, begin to assemble and polymerize. This fibrous corona, which appears as an expanded kinetochore with a crescent or ring shape, acts as a platform that amplifies the SAC signal and facilitates microtubule capture and the conversion of lateral attachments to end-on attachments. MPS1 kinase phosphorylates the dynein adaptor Spindly, causing a conformational change that promotes RZZ polymerization (39), a process crucial for the initial assembly of the fibrous corona. In our experimental setting, treatment with the MPS1 inhibitor Reversine at the final two hours of nocodazole treatment did not affect the BubR1 acetylation (Fig. 4A), while AurkB inhibitors significantly interfered with BubR1 acetylation and MCC maintenance (Fig. 4A-C).

In the previous study in mice, we have shown that BubR1 acetylation is required for MCC maintenance but not the initial assembly, as Mad1 and Mad2 localization to kinetochores were not altered in K243R/+ acetylation-deficient MEFs, while the MCC maintenance was profoundly affected even in the presence of MG132 (15). That inhibition of AurkB kinase activity interferes with AcK250-BubR1 expansion suggest that AurkB activity and BubR1 acetylation appear to function in the maintenance of, rather than the initial establishment of, the fibrous corona and the switch from lateral to end-on attachment. This speculation is supported by the fact that AcK250-BubR1-supported ZW10 expansion, which was abolished upon CENP-E depletion (Fig. 3D). Altogether, AurkB phosphorylation of S16 and S39 enables K250 acetylation of BubR1, which is critically required for fibrous corona maintenance, supporting the conversion of lateral to end-on attachments, synchronized with MCC maintenance. These data are consistent with and extend previous reports that AurkB retards end-on conversion and that BubR1-associated PP2A plays a significant role in end-on conversion, suggesting that the AurkB-BubR1 pathway controls the lateral-to-end-on conversion (53). We have previously demonstrated that AcK250-BubR1 recruits PP2A-B56α, counteracting excessive Aurora B activity following successful KT-MT attachment during chromosome segregation (15). However, we do not rule out the possibility that MPS1 affects AcK250-BubR1 expansion at the kinetochore, as MPS1 is required for polymerization of fibrous corona. It is speculated that MPS1 is crucial for initial establishment of fibrous corona, while AurkB-BubR1-AcK250 is required for the maturation or the sustain of the polymerized corona. Nevertheless, our findings suggest that AurkB-BubR1-AcK250 acetylation controls the switch from lateral to end-on attachment in coordination with the maintenance of the fibrous corona and SAC activity.

At this stage, the mechanism by which two serine residues within the TPR domain affect K250 acetylation remains unclear. Since S39 meets the canonical substrate requirements of AurkB while S16 does not, it is possible that S39 is the primary site and that S16 evolved as an additional regulatory site. Nevertheless, phosphorylation at these sites may induce structural changes that expose K250 for acetylation. Cryo-EM structural analysis of the APC/C-MCC complex (68) indicates that S39 is positioned adjacent to the Mad2-binding site but does not interfere with Mad2 or MCC formation (Supplementary Fig. S6B). Therefore, it is speculated that S39 and S16 phosphorylation of BubR1 at the TPR domain is coordinated with the overall structure of the kinetochore.

In summary, we have delineated a sequential phosphorylation-acetylation SAC signaling code converging on BubR1, linking KT-MT attachment status with control of the lateral-to-end-on attachment switch, and synchronizing these events with maintenance of the fibrous corona and MCC. Given that the BRCA2 tumor suppressor is involved in BubR1 acetylation and deacetylation (12,34), problems in the AurkB-AcK250-BubR1 pathway is likely associated with tumorigenesis. Therefore, the development of drugs targeting this pathway may have clinical implications for treating cancers of chromosome instability, particularly BRCAness cancers.

## Supporting information

time-lapse video

## DATA AVAILABILITY

This paper does not include any original code. All software tools used are described in the Methods section.

Additional information required to reanalyze the data presented in this study can be obtained from the lead contact upon request.

## SUPPLEMENTARY DATA

Supplementary Data are available at NAR online.

## AUTHOR CONTRIBUTIONS

S.C. conceived, performed major experiments, analyzed data, and wrote the manuscript. H.K. & S-S.K. carried out mass spectrometry. H.K. and S.P. generated alanine substitution variants. J.L. conducted some immunofluorescence assays. H.L. conceived, designed, led the study, and wrote the manuscript.

## ACKNOWLEDGMENTS

We extend our gratitude to the members of the Lee Lab for their invaluable discussions and insights, which greatly contributed to this study. Haemin Park helped dealing with radioisotopes in the *in vitro* phosphorylation assays. The Structured Illumination Microscopy (SIM) images were acquired using the AP DeltaVision OMX Ultra High-Resolution Fluorescence Microscope at the SNU Center for Macromolecular and Cell Imaging (CMCI) in IMBG. The graphical abstract was designed by Soyoung Joo. The mass spectrometry proteomics data have been deposited to the ProteomeXchange Consortium via the PRIDE partner repository with the dataset identifier PXD066850 and 10.6019/PXD066850 (69).

