## Supplementary Figures and Table for "Linking Kinetochore Attachment to Checkpoint Control: The Role of Aurora B in BubR1 Acetylation"

### Figure S1

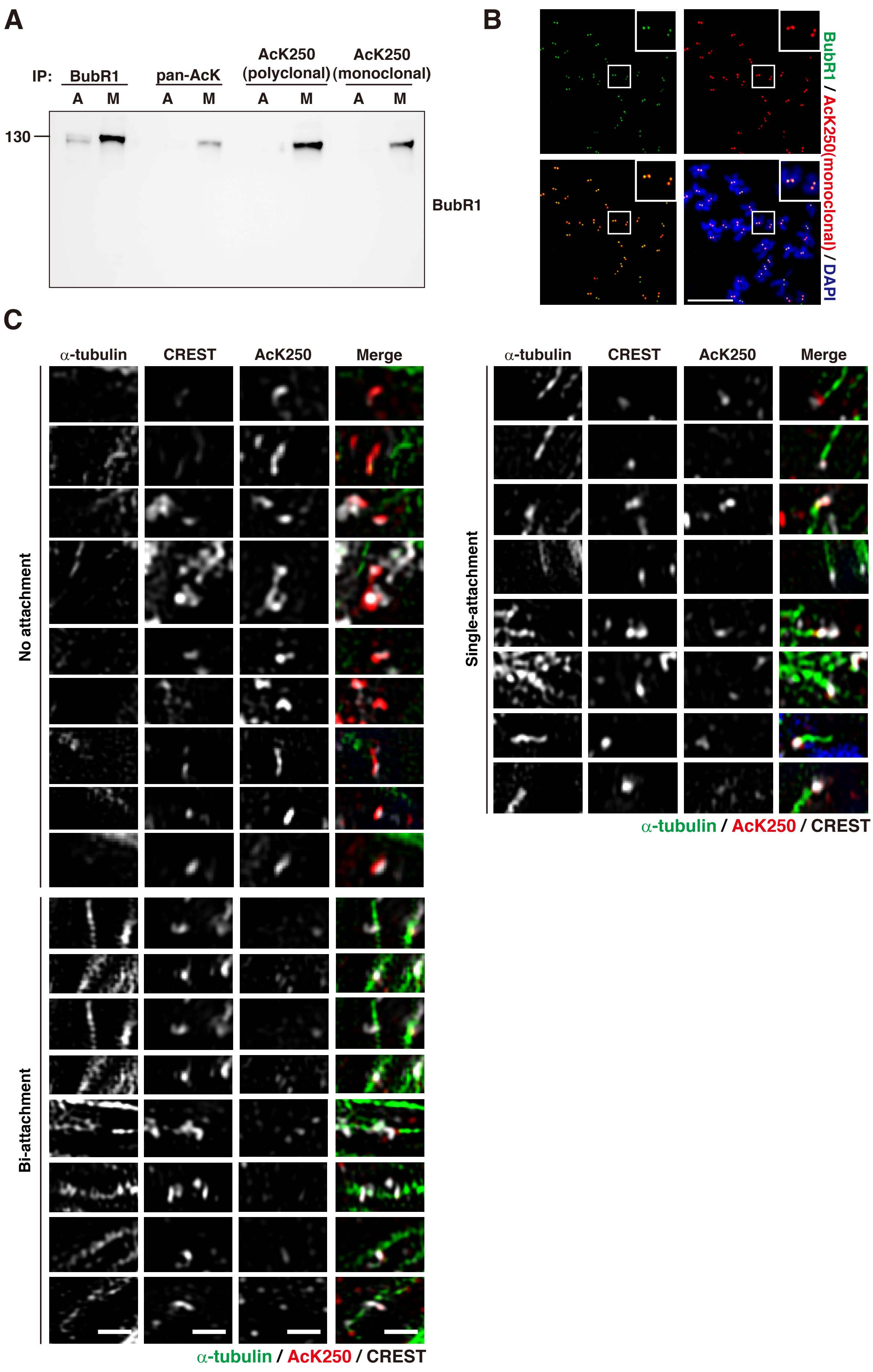
A
IP:
BubR1
pan-AcK
AcK250 (polyclonal)
AcK250 (monoclonal)
A M A M A M A M
130
BubR1
B
BubR1 / AcK250(monoclonal) / DAPI
C
α-tubulin
CREST
AcK250
Merge
No attachment
Bi-attachment
Single-attachment
α-tubulin / AcK250 / CREST
α-tubulin / AcK250 / CREST

### Figure S2

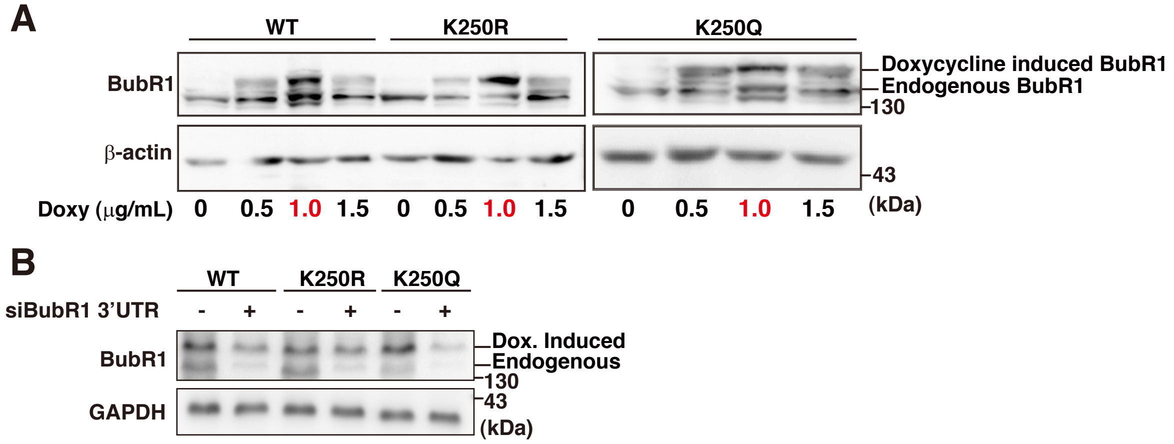
A
WT
K250R
K250Q
BubR1
β-actin
Doxycycline induced BubR1
Endogenous BubR1
130
43
(kDa)
Doxy (μg/mL) 0 0.5 1.0 1.5 0 0.5 1.0 1.5 0 0.5 1.0 1.5
B
WT
K250R
K250Q
siBubR1 3'UTR - + - + - +
BubR1
GAPDH
Dox. Induced
Endogenous
130
43
(kDa)

### Figure S3

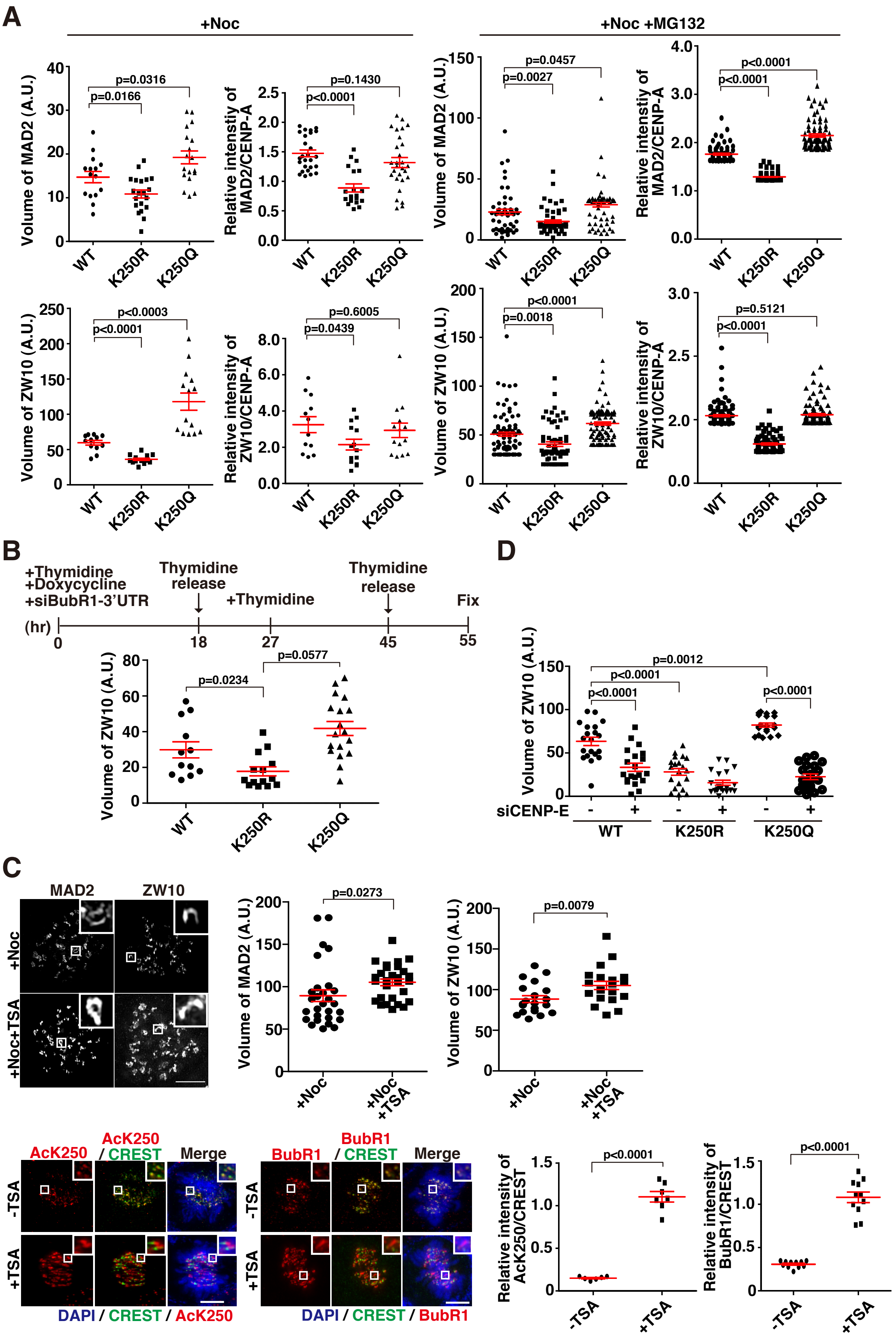
A
+Noc
+Noc +MG132
Volume of MAD2 (A.U.)
Relative intenstiy of MAD2/CENP-A
Volume of ZW10 (A.U.)
Relative intensity of ZW10/CENP-A
WT
K250R
K250Q
p=0.0316
p=0.0166
p=0.1430
p<0.0001
p=0.0457
p=0.0027
p<0.0001
p<0.0001
p<0.0003
p<0.0001
p=0.6005
p=0.0439
p<0.0001
p=0.0018
p=0.5121
p<0.0001
B
+Thymidine
+Doxycycline
+siBubR1-3'UTR
Thymidine release
+Thymidine
Thymidine release
Fix
(hr)
0
18
27
45
55
p=0.0234
p=0.0577
D
p=0.0012
p<0.0001
p<0.0001
p<0.0001
siCENP-E
C
MAD2
ZW10
+Noc
+Noc+TSA
p=0.0273
p=0.0079
AcK250
AcK250 / CREST
Merge
-TSA
+TSA
DAPI / CREST / AcK250
BubR1
BubR1 / CREST
DAPI / CREST / BubR1
Relative intensity of AcK250/CREST
Relative intensity of BubR1/CREST
p<0.0001
p<0.0001

### Figure S4

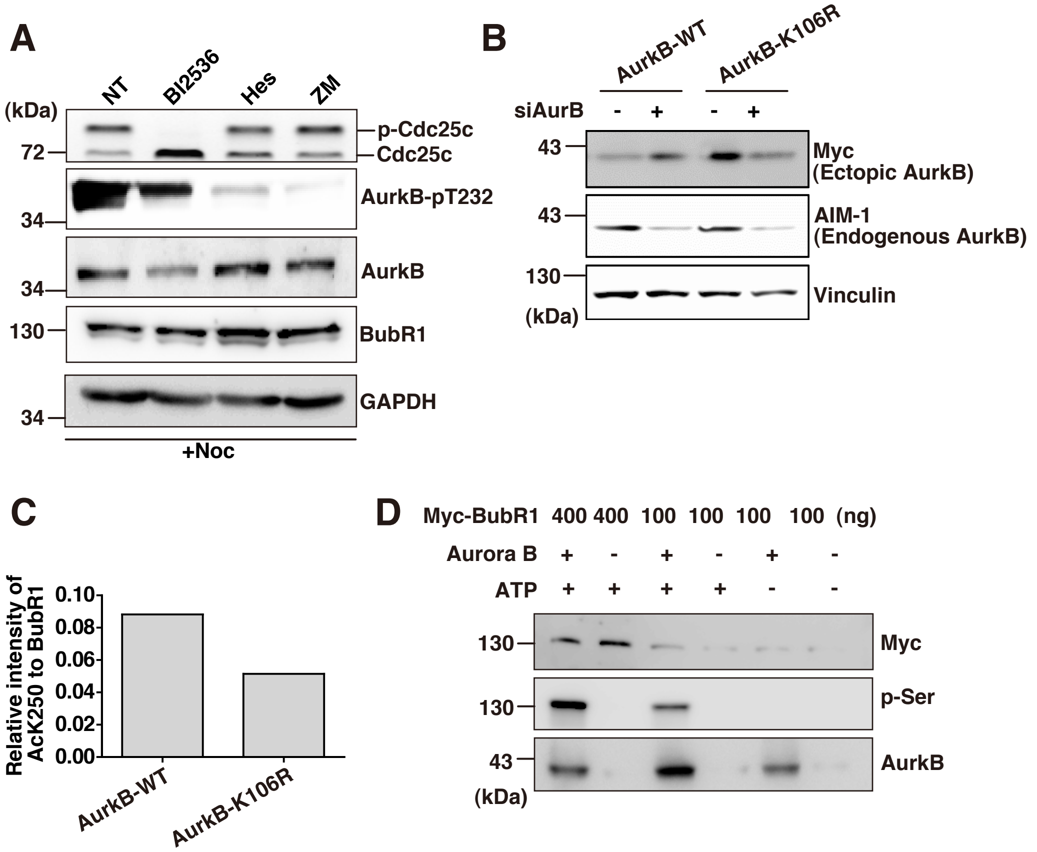
A
NT
BI2536
Hes
ZM
(kDa)
72
34
34
130
34
p-Cdc25c
Cdc25c
AurkB-pT232
AurkB
BubR1
GAPDH
+Noc
B
AurkB-WT
AurkB-K106R
siAurB
- + - +
43
43
130
(kDa)
Myc
(Ectopic AurkB)
AIM-1
(Endogenous AurkB)
Vinculin
C
Relative intensity of AcK250 to BubR1
0.10
0.08
0.06
0.04
0.02
0.00
AurkB-WT
AurkB-K106R
D
Myc-BubR1 400 400 100 100 100 100 (ng)
Aurora B + - + - + -
ATP + + + + - -
130
130
43
(kDa)
Myc
p-Ser
AurkB

### Figure S5

**A**

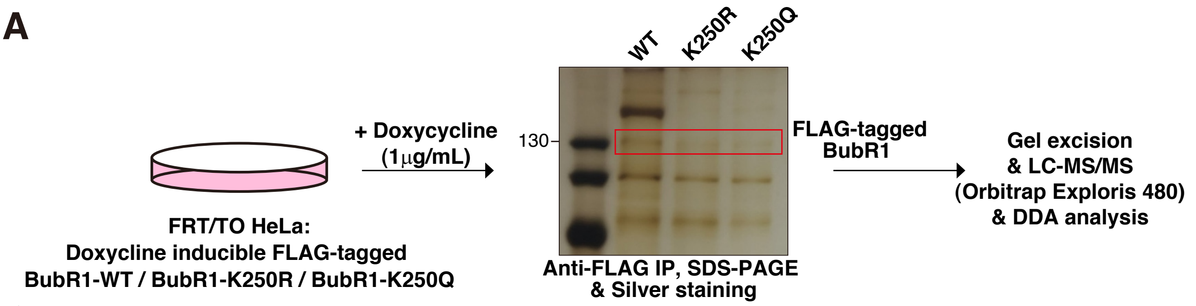

| Site / Genotype | S16 | S39 | S49 | S361 | S411 | T600 | S619 | S649 | T654 | T493 S495 | T508 S509 | T434 S435 |
|---|---|---|---|---|---|---|---|---|---|---|---|---|
| WT | -P | -P | - | -P | -P | -P | -P | -P | -P | -P | -P | -P |
| K250R | - | - | - | -P | - | - | -P | - | -P | -P | -P | - |
| K250Q | -P | -P | -P | -P | -P | -P | -P | -P | -P | - | -P | -P |

**B**

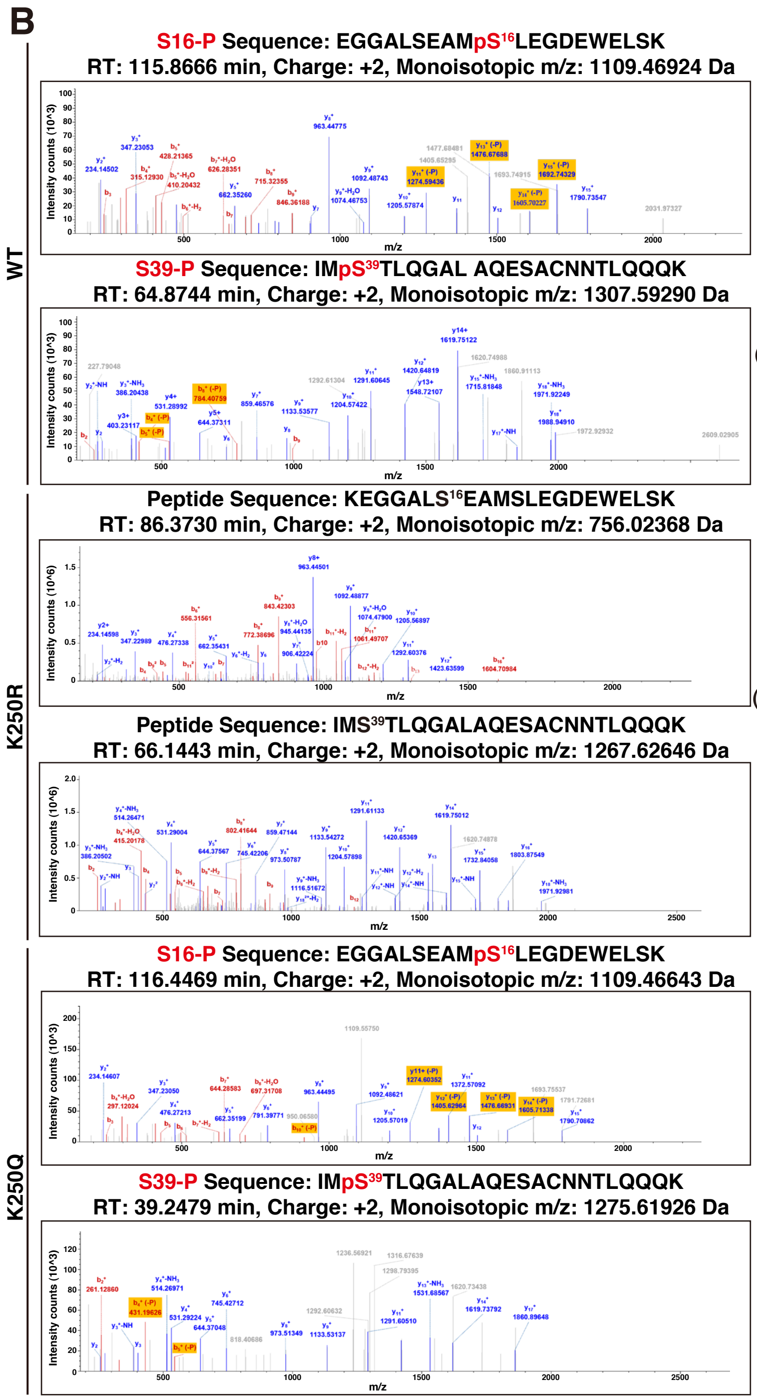

**C**

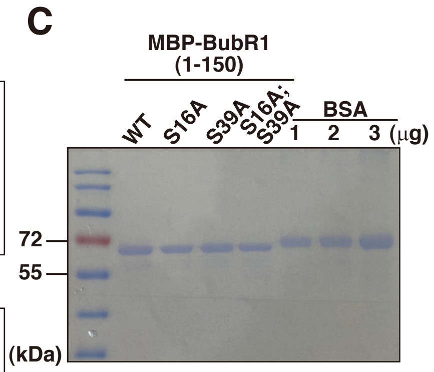

**D**

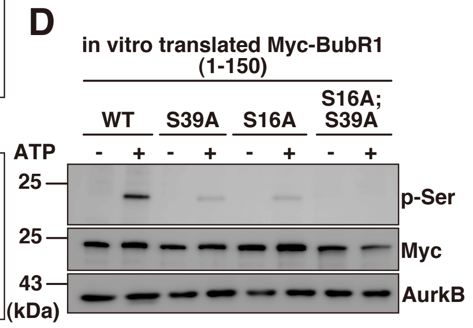

### Figure S6

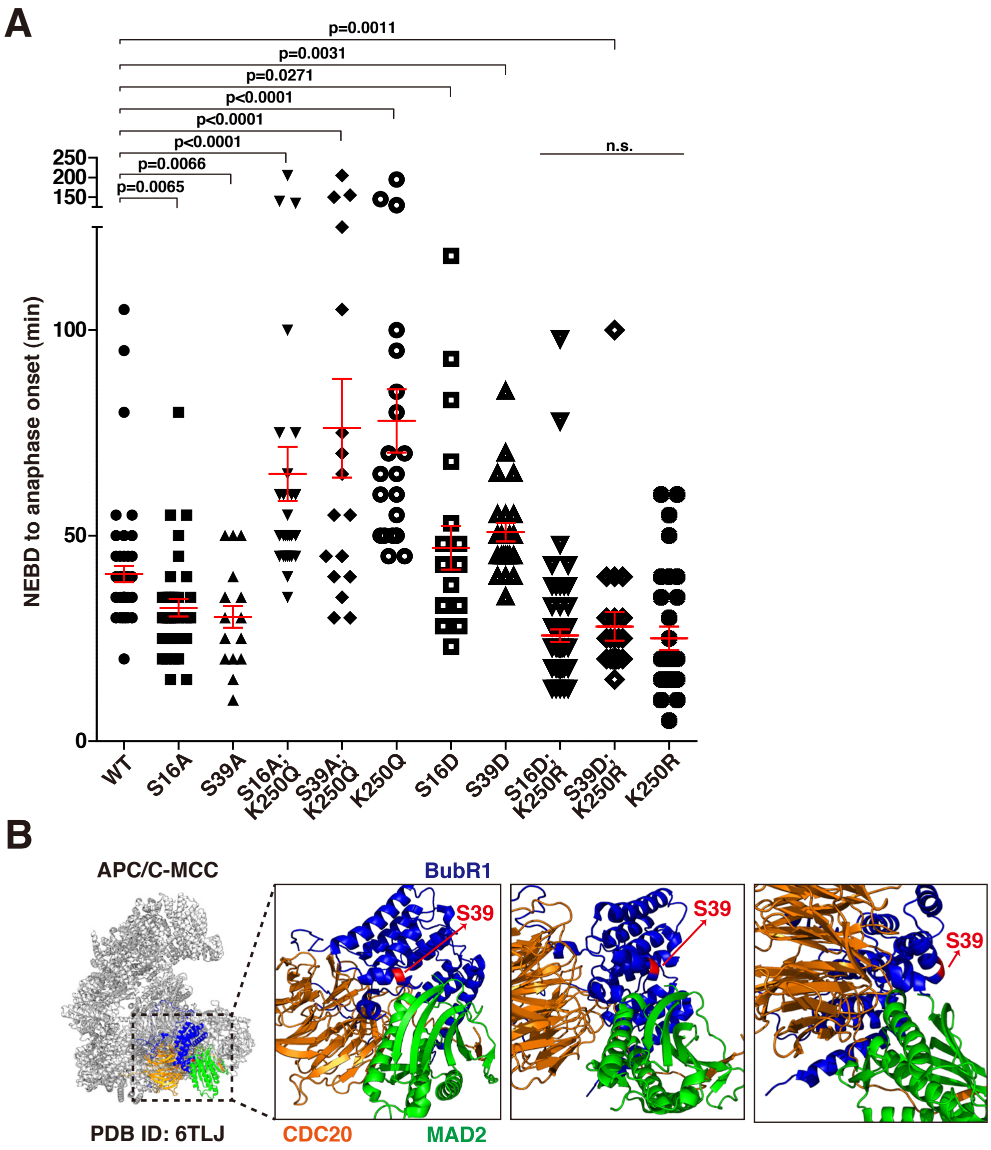
A
p=0.0011
p=0.0031
p=0.0271
p<0.0001
p<0.0001
p<0.0001
p=0.0066
p=0.0065
n.s.
NEBD to anaphase onset (min)
250
200
150
100
50
0
WT
S16A
S39A
S16A; K250Q
S39A; K250Q
K250Q
S16D
S39D
S16D; K250R
S39D; K250R
K250R
B
APC/C-MCC
PDB ID: 6TLJ
BubR1
S39
S39
S39
CDC20
MAD2

**Table S1. Identification of BubR1 phosphorylation sites upon unattachment using LC-MS/MS analysis (related to Figure 5).**

Results from three independent LC-MS/MS experiments. Sequence coverage for BubR1-WT, -K250R, -K250Q was 73.85%, 66.62%, and 66.62%, respectively. Phosphorylation published already are indicated in bold.

| | WT 1 | WT 2 | WT 3 | K250R 1 | K250R 2 | K250R 3 | K250Q 1 | K250Q 2 | K250Q 3 | **Kinase** | **Reference** |
|---|---|---|---|---|---|---|---|---|---|---|---|
| S12 | | | | -P | | | | | | | |
| S16 | -P | | | | | | -P | | | | |
| S39 | -P | -P | -P | | | | | -P | -P | | |
| T40 | -P | -P | -P | -P | -P | | | -P | -P | | |
| S49 | | | | | | | | -P | -P | | |
| **T54** | | | | | | | | | | - | 1 |
| Y81 | -P | | | | | | | | | | |
| S83 | -P | -P | | -P | | | | | | | |
| T204 | | -P | -P | | -P | | | -P | -P | | |
| S361 | | | -P | -P | | -P | | | -P | | |
| **S367** | | | | | | | | | | - | 2 |
| **S384** | | | | | | | | | | - | 3 |
| Y404 | | | | -P | | | | | | | |
| S411 | | -P | -P | | | | | -P | | | |
| **T434** | | -P | | | | | | -P | | - | 3,4 |
| **S435** | | -P | -P | | | | | -P | -P | Mps1 (partial)[5] | 2-6 |
| **S453** | | | | | | | | | | - | 5 |
| **T471** | | | | | | | | -P | -P | - | 7 |
| T493 | | | -P | -P | -P | -P | | | | | |
| S495 | | | -P | -P | -P | -P | | | | | |
| S496 | -P | | | | | | | | | | |
| T508 | -P | | -P | -P | | -P | | | | | |
| **S509** | -P | -P | -P | -P | | -P | | | | - | 3 |
| **S543** | | -P | -P | | | | | -P | | Mps1 (partial)[5] | 2-5,7,8 |
| **S564** | | | | | -P | | | | | - | 3 |
| **S574** | | | | | | | | | | - | 3,8 |
| S599 | | -P | | | -P | | | -P | | | |
| Y600 | -P | | -P | | | | | | -P | | |
| **T608** | -P | | | -P | | | -P | | | - | 9 |
| S619 | -P | | | -P | | -P | -P | | | | |
| **T620** | | -P | | -P | -P | | | -P | -P | Cdk1[10] | 10,11 |

| | | | | | | | | | | | |
|---|---|---|---|---|---|---|---|---|---|---|---|
| T648 | -P | | -P | -P | | | -P | | -P | | |
| S649 | -P | -P | | | | | -P | -P | | | |
| T654 | -P | -P | -P | -P | -P | -P | -P | -P | -P | | |
| T658 | | -P | -P | -P | | | -P | | | | |
| **Y660** | | | | -P | | | -P | | | - | 3 |
| S661 | | | | -P | | | -P | | | | |
| **S665** | -P | | | -P | -P | -P | | | | - | 2 |
| **S670** | -P | -P | -P | | -P | -P | -P | -P | -P | Mps1 (partial)[5] | 2-8,12 |
| **S676** | -P | -P | -P | | -P | | | -P | | Plk1[10] | 3,5,10 |
| **T680** | | | | | | | | -P | -P | Plk1[13] | 3,13 |
| S682 | | -P | -P | | | | | -P | -P | | |
| S686 | | -P | -P | | | -P | | -P | -P | | |
| S688 | | | | | | | | -P | -P | | |
| **S697** | -P | | | | | | | | | - | 2 |
| **T710** | -P | | | -P | -P | | -P | | | - | 3 |
| T713 | -P | | | -P | -P | | -P | -P | -P | | |
| **S714** | -P | | | -P | -P | | -P | | | - | 3 |
| T718 | -P | | | -P | -P | | -P | | | | |
| **S720** | -P | -P | -P | -P | -P | -P | -P | -P | -P | - | 3,8 |
| S724 | | | | -P | -P | | -P | | | | |
| Y766 | | | | | | | | -P | | | |
| Y772 | | | | | | | | -P | -P | | |
| **T792** | | | | | | | | | | Plk1[14] | 5,14 |
| T798 | | | | | | | | -P | -P | | |
| **S884** | | -P | | | | | | -P | -P | - | 3 |
| T926 | | -P | -P | | -P | | | | | | |
| S928 | | | | | | | | -P | -P | | |
| S976 | | | | | -P | | | | | | |
| S981 | -P | -P | -P | -P | | -P | -P | | | | |
| S985 | | | | -P | -P | -P | | -P | -P | | |
| **T1008** | | | | | | | | | | Plk1[14] | 5,14 |
| **S1043** | | | | | -P | -P | | | | Mps1 (partial)[5] | 3-5,7,8 |
| T1052 | | | | | -P | | | | | | |
| S1056 | | | | | -P | -P | | | | | |
| T1057 | | | | | -P | | | | | | |
| S1058 | | -P | -P | | -P | -P | | -P | -P | | |
| T1072 | | -P | -P | | | | | -P | -P | | |
| S1091 | | | -P | | | | | -P | -P | | |
| T1122 | | | | | -P | | | | | | |
| T1125 | | | | | -P | | | | | | |

| | | | | | | | | | |
|---|---|---|---|---|---|---|---|---|---|
| T1160 | | | | | | | | -P | -P |
| Y1206 | | -P | -P | | | -P | | -P | -P |
| Y1208 | | -P | -P | | | -P | | -P | -P |
| T1216 | | -P | -P | | -P | | | | |
| T1249 | | -P | -P | | | | | -P | -P |
| Y1263 | | | | | | | | -P | -P |

1. Imami, K., Sugiyama, N., Kyono, Y., Tomita, M., and Ishihama, Y. (2008). Automated Phosphoproteome Analysis for Cultured Cancer Cells by Two-Dimensional NanoLC-MS Using a Calcined Titania/C18 Biphasic Column. Analytical Sciences *24*, 161-166. 10.2116/analsci.24.161.
2. Zhou, H., Di Palma, S., Preisinger, C., Peng, M., Polat, A.N., Heck, A.J., and Mohammed, S. (2013). Toward a comprehensive characterization of a human cancer cell phosphoproteome. J Proteome Res *12*, 260-271. 10.1021/pr300630k.
3. Ochoa, D., Jarnuczak, A.F., Viéitez, C., Gehre, M., Soucheray, M., Mateus, A., Kleefeldt, A.A., Hill, A., Garcia-Alonso, L., Stein, F., et al. (2020). The functional landscape of the human phosphoproteome. Nature Biotechnology *38*, 365-373. 10.1038/s41587-019-0344-3.
4. Olsen, J.V., Vermeulen, M., Santamaria, A., Kumar, C., Miller, M.L., Jensen, L.J., Gnad, F., Cox, J., Jensen, T.S., Nigg, E.A., et al. (2010). Quantitative Phosphoproteomics Reveals Widespread Full Phosphorylation Site Occupancy During Mitosis. Science Signaling *3*, ra3-ra3. 10.1126/scisignal.2000475.
5. Huang, H., Hittle, J., Zappacosta, F., Annan, R.S., Hershko, A., and Yen, T.J. (2008). Phosphorylation sites in BubR1 that regulate kinetochore attachment, tension, and mitotic exit. Journal of Cell Biology *183*, 667-680. 10.1083/jcb.200805163.
6. Dephoure, N., Zhou, C., Villén, J., Beausoleil, S.A., Bakalarski, C.E., Elledge, S.J., and Gygi, S.P. (2008). A quantitative atlas of mitotic phosphorylation. Proceedings of the National Academy of Sciences *105*, 10762-10767. 10.1073/pnas.0805139105.
7. Rigbolt, K.T.G., Prokhorova, T.A., Akimov, V., Henningsen, J., Johansen, P.T., Kratchmarova, I., Kassem, M., Mann, M., Olsen, J.V., and Blagoev, B. (2011). System-Wide Temporal Characterization of the Proteome and Phosphoproteome of Human Embryonic Stem Cell Differentiation. Science Signaling *4*, rs3-rs3. 10.1126/scisignal.2001570.
8. Elowe, S., Dulla, K., Uldschmid, A., Li, X., Dou, Z., and Nigg, E. (2010). Uncoupling of the spindle-checkpoint and chromosome-congression functions of BubR1. Journal of cell science *123*, 84-94. 10.1242/jcs.056507.
9. Guo, Y., Kim, C., Ahmad, S., Zhang, J., and Mao, Y. (2012). CENP-E--dependent BubR1 autophosphorylation enhances chromosome alignment and the mitotic checkpoint. J Cell Biol *198*, 205-217. 10.1083/jcb.201202152.
10. Elowe, S., Hümmer, S., Uldschmid, A., Li, X., and Nigg, E.A. (2007). Tension-sensitive Plk1 phosphorylation on BubR1 regulates the stability of kinetochore microtubule interactions. Genes Dev *21*, 2205-2219. 10.1101/gad.436007.
11. Cordeiro, M.H., Smith, R.J., and Saurin, A.T. (2020). Kinetochore phosphatases suppress autonomous Polo-like kinase 1 activity to control the mitotic checkpoint. J Cell Biol *219*. 10.1083/jcb.202002020.
12. Daub, H., Olsen, J.V., Bairlein, M., Gnad, F., Oppermann, F.S., Körner, R., Greff, Z., Kéri, G., Stemmann, O., and Mann, M. (2008). Kinase-selective enrichment enables quantitative phosphoproteomics of the kinome across the cell cycle. Mol Cell *31*, 438-448. 10.1016/j.molcel.2008.07.007.
13. Suijkerbuijk, Saskia J.E., Vleugel, M., Teixeira, A., and Kops, Geert J.P.L. (2012). Integration of Kinase and Phosphatase Activities by BUBR1 Ensures Formation of Stable Kinetochore-Microtubule Attachments. Developmental Cell *23*, 745-755. https://doi.org/10.1016/j.devcel.2012.09.005.
14. Matsumura, S., Toyoshima, F., and Nishida, E. (2007). Polo-like kinase 1 facilitates chromosome alignment during prometaphase through BubR1. J Biol Chem *282*, 15217-15227. 10.1074/jbc.M611053200.

**Supplemental Figure Legends**

**Figure S1. Three-dimensional structured illumination microscopy (SIM) images of kinetochores with different attachment states.** (A) Assessing the specificity of the monoclonal antibody to K250 acetylated BubR1 (AcK250) in Nocodazole-arrested and mitotic shake-off HeLa cells (M). IP with the polyclonal antibody to AcK250-BubR1 (11) or rabbit monoclonal Ack250-BubR1 were performed, followed by WB with anti-BubR1 antibody (BD Biosciences). To compare, IP with anti-BubR1 or anti-pan-Acetyl-lysine antibody were employed. Attached cells (A) were employed for comparison. (B) Assessing the specificity of anti-AcK250 mAb in immunofluorescence. Metaphase chromosome spreads from HeLa cells were subjected to co-immunostaining with anti-BubR1 (Green) and anti-AcK250 mAb (Red). For metaphase chromosome spread, HeLa cells were treated with 100ng/ml of Colcemid for 20 h. (C) The SIM immunofluorescence images of various states of mitotic kinetochores. HeLa cells were treated with 10μM of MG132 and subjected to co-immunostaining of acetylated BubR1 (AcK250 mAb, Red), CREST (White), and anti-α-tubulin (Green) antibodies. CREST-immunostained kinetochores were grouped in three different attachment states: No attachment (No); monotelic (Single-attachment); amphitelic (Bi-attachment). Note that there is no AcK250 at the kinetochore in amphitelic attachment. AcK250-positive immunofluorescence is the highest in No attachment.

**Figure S2. Efficiency of the 3'UTR-targeting siBubR1 in depleting endogenous BubR1 in doxycycline-inducible HeLa FRT-TO BubR1 cell lines.** (A) Increasing doses of doxycycline were treated in HeLa-FRT-TO cells to induce single-copy expression of *BubR1-WT*, *BubR1-K250R*, *BubR1-K250Q*, respectively(1-3). WB with anti-BubR1 antibody was performed 48 hours post induction of BubR1 expression with indicated doxycycline treatment. Same blot was reprobed with anti-β-actin antibody for loading control. (B) Depletion of endogenous

BubR1 in doxycycline-treated FRT/TO cells after siBubR1 transfection. Cells were treated with 3'UTR-targeting siBubR1 and doxycycline simultaneously for 48 hours before analysis.

**Figure S3. Measuring the volume of Mad2 and ZW10 in HeLa-FRT-TO cells expressing *BubR1-WT*, *-K250R*, or *-K250Q*, and the effect of HDAC inhibitor in fibrous corona expansion.** (A) Relative intensities and volumes of MAD2 and ZW10 at kinetochores in HeLa-FRT-TO cells expressing *BubR1-WT*, *-K250R*, or *-K250Q*, following treatment with nocodazole alone or in combination with MG312. For nocodazole-only treatment: Mad2-*WT*, n=26; *K250R*, n=20; *K250Q*, n=28 and ZW10-*WT*, n=12; *K250R*, n=14; *K250Q*, n=14. For nocodazole + MG132 treatment: Mad2-*WT*, n=69; *K250R*, n=94; *K250Q*, n=76 and ZW10-*WT*, n=111; *K250R*, n=68; *K250Q*, n=98. Fluorescence intensities in SIM images were normalized to co-localized CENP-A signals. Measuring the volume by 3D Objects Counter plugin is described in Materials and Methods (mean±s.e.m.). (B) Measuring the volume of ZW10 following double-thymidine block and release (related to Fig. 3B). HeLa-FRT-TO BubR1-expressing cells were treated with doxycycline for 55 hours to induce the expression of *BubR1-WT*, *-K250R*, or *-K250Q*. Endogenous BubR1 was depleted using siRNA targeting the 3'UTR. Cells were fixed for immunostaining 10 hours post final thymidine release to enrich for mitotic populations. Number of cells scored: *WT*, n=11; *K250R*, n=14; *K250Q*, n=18 (mean±s.e.m.).

(C) (Top) Effects of trichostatin A (TSA), pan-HDAC inhibitor (HDACi), in fibrous corona expansion in nocodazole-arrested cells. Volumes of MAD2 (Noc, n=28; Noc+TSA, n=28) and ZW10 (Noc, n=20; Noc+TSA, n=20) were measured in HeLa cells treated with nocodazole (200ng/mL, 20h) and DMSO (+Noc) or with 10μM of TSA (+Noc+TSA). TSA was added during the final 2 hours of Nocodazole teatment. Subsequently, cells were subjected to immunostaining with anti-MAD2 or anti-ZW10 antibodies (mean±s.e.m.). (Bottom) Effects of

TSA in K250-BubR1 acetylation and stability at the kinetochore. HeLa cells were treated with TSA (10μM) for 6 hours and immunostained with the antibodies indicated. Number of cells scored for the graph at the right: -TSA, n=11; +TSA, n=11 (mean±s.e.m.). (D) Quantification of ZW10 volume in HeLa-FRT-TO cells expressing *BubR1-WT*, *-K250R*, *-K250Q* with or without CENP-E depletion (related to Fig. 3D). Cells were transfected with either control siRNA (siLuciferase, -) or siRNA targeting CENP-E (siCENP-E, +). Number of cells scored: *WT*, siCtrl, n=20; *WT*, siCENP-E, n=19; *K250R*, siCtrl, n=20; *K250R*, siCENP-E, n=20; *K250Q*, siCtrl, n=21; *K250Q*, siCENP-E, n=18 (mean±s.e.m.).

**Figure S4. Validation of various kinase inhibitor in AurkB activity and AcK250 acetylation.** (A) Western blot analysis validating the efficiency of various kinase inhibitors. The activity of the Plk1 inhibitor BI2536 was assessed by measuring levels of phosphorylated Cdc25C, a known Plk1 substrate. The effects of Aurora B inhibitors were evaluated by detecting Aurora B auto-phosphorylation at threonine 232 (AurkB-pT232). Kinase inhibitors were applied during the final 2 hours of a 20-hour nocodazole treatment. (Related to Fig. 4A and B). (B) Western analysis showing the expression level of wild-type Aurora B (*AurB-WT*), kinase-dead Aurora B (*AurB-K106R*), and depletion of endogenous Aurora B (AIM-1). siRNA targeting endogenous Aurora B was transfected for 48 hours. (C) Quantification of relative AcK250 intensity normalized to BubR1 in cells expressing *AurB-WT* and AurB-*K106R*. (*AurB-WT*: 0.088215, n=41, AurB-*K106R*: 0.051434, n=65) (related Figure 4D and F). (D) *In vitro* phosphorylation assay showing that BubR1 is phosphorylated by Aurora B. Myc-tagged BubR1 was generated via *in vitro* transcription and translation. Either 400 ng or 100 ng of the final product was employed. Recombinant Aurora B (200ng) and ATP (100μM) were employed for phosphorylation. The degree of phosphorylation was detected in the presence of AurkB (p-

Ser). WB with 9E10 (anti-Myc) at the top shows the level of *in vitro* translated BubR1 employed in the assay (Myc), and the WB with anti-AurkB indicates the presence of AurkB employed in the assay (AurkB).

**Figure S5. Overview of the LC/MS-MS workflow and the validation of S16 and S39 as AurkB phosphorylation sites.** (A) Schematic illustration of the LC/MS-MS analysis. FLAG-tagged *BubR1-WT*, *-K250R*, and *-K250Q* were expressed in HeLa-FRT-TO cells through doxycycline (1μg/mL) treatment for 48 hours. Cells were then treated with nocodazole for 20 hours prior to harvest. Lysates from various HeLa-FRT-TO *BubR1*-expressing cells were subjected to IP with anti-FLAG, followed by silver staining. BubR1 bands predicted by size were excised and analyzed by LC-MS/MS with data-dependent acquisition (DDA) to identify phosphorylation sites. The bottom panel compares the LC-MS/MS phosphorylation results in wild-type, *-K250R*, and *-K250Q* expressing HeLa-FRT-TO cells. (B) Mass spectrometry-derived intensity counts of peptide fragments containing serines 16 and 39 from BubR1-WT, -K250R and -K250Q. Retention time (RT), charge state, monoisotopic m/z, and peptide sequences of the detected fragments are indicated. (C) Coomassie Brilliant Blue (CBB) staining to assess the purity and quantity of MBP-tagged BubR1 purified by size exclusion chromatography. The purified peptides were subsequently used for the in vitro phosphorylation assay shown in Fig. 5E. (D) *In vitro* phosphorylation assay using Myc-tagged N-terminally truncated BubR1 (residues 1–150), produced via *in vitro* transcription and translation, and recombinant Aurora B. A total of 400 ng of BubR1 protein was used in the phosphorylation reaction. ATP (100μM) was included in the indicated lanes. Phosphorylation was detected by Western blotting using a phospho-specific antibody.

**Figure S6. Predicted structural positioning of Serine 39 of BubR1 in APC/MCC.** (A) Timing of NEBD to anaphase onset from Figure 6D. Number of cells scored: *WT*, n=55; *S16A*, n=37; *S39A*, n=19; *S16A;K250Q*, n=30; *S39A;K250Q*, n=18; *K250Q*, n=24; *S16D*, n=21; *S39D*, n=24; *S16D;K250R*, n=74; *S39D;K250R*, n=24; *K250R*, n=30. Graphs are the result from three independent experiments (mean±s.e.m.). (B) Cryo-EM structure of the APC/C-MCC complex (66), PDB ID: 6TLJ. Serine 39 (red) of BubR1 is located near the interface between BubR1 (blue) and MAD2 (green) but does not interfere with MAD2 interaction.

**Supplemental Videos**

**Video S1**. Time-lapse live-cell imaging of H2B-GFP-expressing HeLa cells transfected with siBubR1-3'UTR, targeting endogenous BubR1, and mCherry-BubR1-WT, related to Fig. 6D and supplementary figure S6A.

**Video S2**. Time-lapse live-cell imaging of H2B-GFP-expressing HeLa cells transfected with siBubR1-3'UTR, targeting endogenous BubR1, and mCherry-BubR1-S16A, related to Fig. 6D and supplementary figure S6A.

**Video S3**. Time-lapse live-cell imaging of H2B-GFP-expressing HeLa cells transfected with siBubR1-3'UTR, targeting endogenous BubR1, and mCherry-BubR1-S39A, related to Fig. 6D and supplementary figure S6A.

**Video S4**. Time-lapse live-cell imaging of H2B-GFP-expressing HeLa cells transfected with siBubR1-3'UTR, targeting endogenous BubR1, and mCherry-BubR1-S16A;K250Q, related to Fig. 6D and supplementary figure S6A.

**Video S5**. Time-lapse live-cell imaging of H2B-GFP-expressing HeLa cells transfected with siBubR1-3'UTR, targeting endogenous BubR1, and mCherry-BubR1-S39A;K250Q, related to Fig. 6D and supplementary figure S6A.

**Video S6**. Time-lapse live-cell imaging of H2B-GFP-expressing HeLa cells transfected with siBubR1-3'UTR, targeting endogenous BubR1, and mCherry-BubR1-K250Q, related to Fig. 6D and supplementary figure S6A.

**Video S7**. Time-lapse live-cell imaging of H2B-GFP-expressing HeLa cells transfected with siBubR1-3'UTR, targeting endogenous BubR1, and mCherry-BubR1-S16D, related to Fig. 6D and supplementary figure S6A.

**Video S8**. Time-lapse live-cell imaging of H2B-GFP-expressing HeLa cells transfected with siBubR1-3'UTR, targeting endogenous BubR1, and mCherry-BubR1-S39D, related to Fig. 6D and supplementary figure S6A.

**Video S9**. Time-lapse live-cell imaging of H2B-GFP-expressing HeLa cells transfected with siBubR1-3'UTR, targeting endogenous BubR1, and mCherry-BubR1-S16D;K250R, related to Fig. 6D and supplementary figure S6A.

**Video S10**. Time-lapse live-cell imaging of H2B-GFP-expressing HeLa cells transfected with siBubR1-3'UTR, targeting endogenous BubR1, and mCherry-BubR1-S39D;K250R, related to Fig. 6D and supplementary figure S6A.

**Video S11**. Time-lapse live-cell imaging of H2B-GFP-expressing HeLa cells transfected with siBubR1-3'UTR, targeting endogenous BubR1, and mCherry-BubR1-K250R, related to Fig. 6D and supplementary figure S6A.
